# Binding of an X-specific condensin correlates with a reduction in active histone modifications at gene regulatory elements

**DOI:** 10.1101/516419

**Authors:** Lena Annika Street, Ana Karina Morao, Lara Heermans Winterkorn, Chen-Yu Jiao, Sarah Elizabeth Albritton, Mohammed Sadic, Maxwell Kramer, Sevinç Ercan

**Author notes:** Correspondence should be addressed to: Sevinc Ercan.

## Abstract

Condensins are evolutionarily conserved protein complexes that are required for chromosome segregation during cell division and genome organization during interphase. In *C. elegans*,, a specialized condensin, which forms the core of the dosage compensation complex (DCC), binds to and represses X chromosome transcription. Here, we analyzed DCC localization and the effect of DCC depletion on histone modifications, transcription factor binding, and gene expression using ChIP-seq and mRNA-seq. Across the X, DCC accumulates at accessible gene regulatory sites in active chromatin and not heterochromatin. DCC is required for reducing the levels of activating histone modifications, including H3K4me3 and H3K27ac, but not repressive modification H3K9me3. In X-to-autosome fusion chromosomes, DCC spreading into the autosomal sequences locally reduces gene expression, thus establishing a direct link between DCC binding and repression. Together, our results indicate that DCC-mediated transcription repression is associated with a reduction in the activity of X chromosomal gene regulatory elements.

**SUMMARY:** Condensins are evolutionarily conserved protein complexes that mediate chromosome condensation during cell division and have been implicated in gene regulation during interphase. Here, we analyzed the gene regulatory role of an X-specific condensin (DCC) in *C. elegans*, by measuring its effect on histone modifications associated with transcription regulation. We found that in X-to-autosome fusion chromosomes, DCC spreading into autosomal sequences locally reduces gene expression, establishing a direct link between DCC binding and repression. DCC is required for reduced levels of histone modifications associated with transcription activation at X chromosomal promoters and enhancers. These results are consistent with a model whereby DCC binding directly or indirectly results in a reduction in the activity of X chromosomal gene regulatory elements through specific activating histone modifications.

## BACKGROUND

Regulation of chromosome structure is essential for the establishment and maintenance of accurate gene expression. A key regulator of chromosome structure across all organisms is condensin, a multi-subunit protein complex that belongs to the family of <u>s</u>tructural <u>m</u>aintenance of <u>c</u>hromosomes (SMC) complexes (Hirano 2006; van Ruiten and Rowland 2018). Condensins are required for chromosome condensation and segregation in all eukaryotes (Hirano 2016). Condensins are also important for genome organization and have been implicated in gene regulation during interphase (Paul *et al*., 2018). Genome-wide binding experiments indicate that condensins bind to a subset of gene regulatory elements including promoters, enhancers, tRNA genes, and topologically associated domain (TAD) boundaries (Jeppsson *et al*., 2014). However, the link between condensin binding at these sites and its function in gene regulation remains unknown.

Here we addressed the link between condensin and transcription by using a clear paradigm for the gene-regulatory function of condensins, the *C. elegans*, <u>d</u>osage <u>c</u>ompensation complex (DCC). Like most metazoans, *C. elegans*, contain two types of condensins (I and II) that partially differ in their subunit composition, chromosomal binding and function (Csankovszki *et al*., 2009). in addition to the canonical condensins, *C. elegans*, contain a third condensin, condensin I^DC^ (hereafter DC), that differs from condensin i by a single SMC-4 variant, DPY-27 (Csankovszki *et al*., 2009). Condensin DC interacts with additional subunits necessary for DCC binding and function (Figure 1A) (reviewed in (Albritton and Ercan 2017)). The DCC specifically binds to both hermaphrodite X chromosomes and represses each by half to equalize X chromosomal transcript levels between XX hermaphrodites and XO males (Jans *et al*., 2009; Kruesi *et al*., 2013; Kramer *et al*., 2015; Kramer *et al*., 2016).

**Figure 1.**
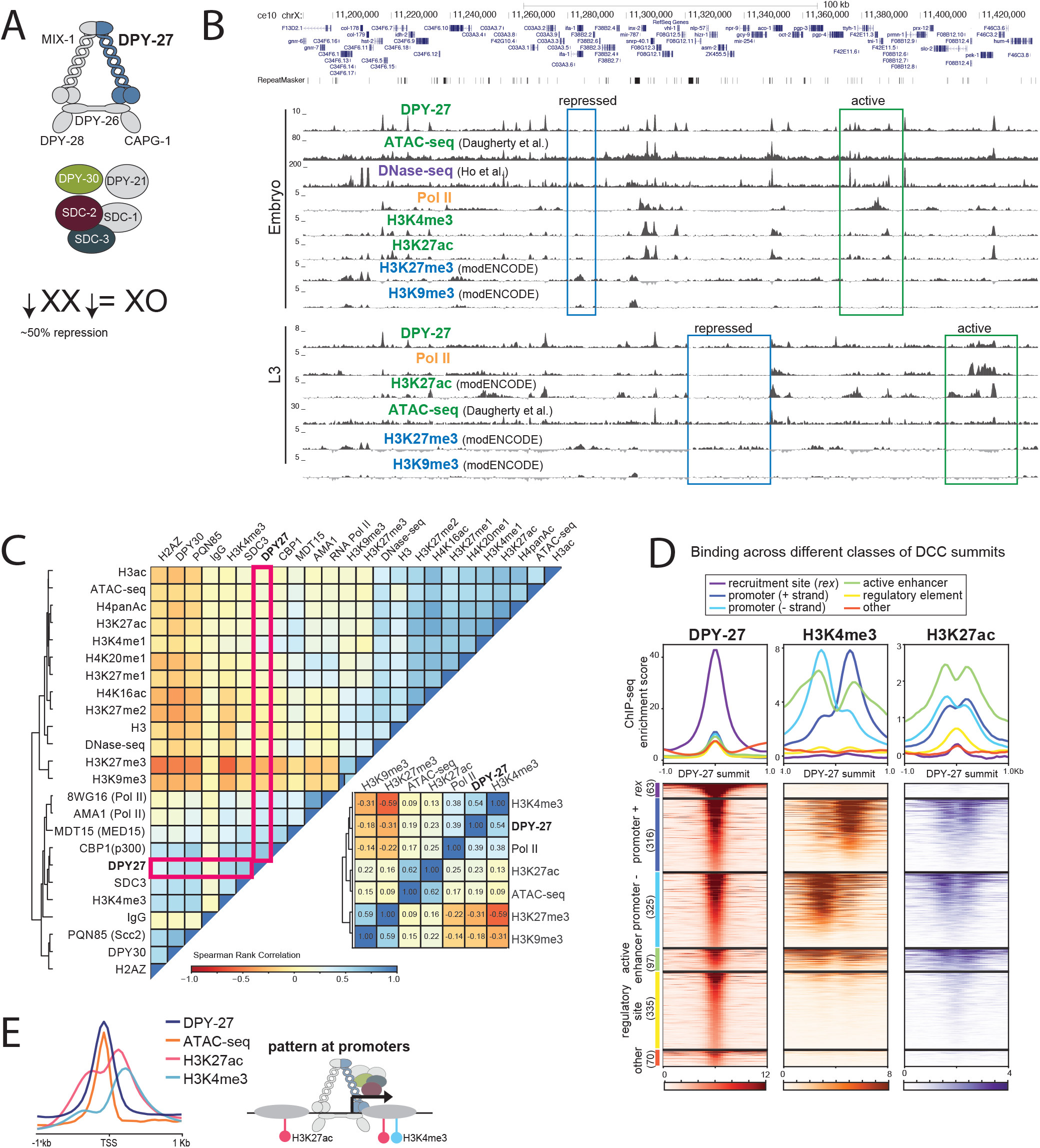
DCC binding correlates with active chromatin marks at gene regulatory elements. **(A)** The *C. elegans*, dosage compensation complex (DCC) contains a specialized condensin complex (condensin DC) that is distinguished from canonical condensin I by a single SMC-4 variant, DPY-27. The non-condensin DCC subunits SDC-2, SDC-3, and DPY-30 interact with condensin DC, and are required for its recruitment to the X chromosomes. DPY-21 is a histone demethylase that converts H4K20me2 to H4K20me1. DCC binds to and represses X chromosomes in hermaphrodites by approximately two-fold. **(B)** ChIP-seq, DNase-seq (Ho *et al*., 2017) and ATAC-seq (Daugherty *et al*., 2017) profiles at a representative 250kb region of the X chromosome in embryos and L3 larval stage worms. Example active and repressed chromatin regions are labeled in green and blue, respectively. DPY-27 (DCC) binding overlaps with Pol II binding, active chromatin marks, and accessible regions (ATAC-seq). **(C)** Spearman rank correlation values are shown for average ChIP- seq scores of histone modifications, ATAC-seq and DNase-seq signals within 1kb contiguous windows across the X chromosome in wild type embryos. Zoomed in plot highlights that DCC (DPY-27) binding positively-correlates more with promoter marks (H3K4me3) and Pol II, with active enhancers (H3K27ac) and regulatory regions (ATAC-seq), and negatively-correlates with repressive marks (H3K27me3, H3K9me3). **(D)** The DPY-27 ChIP-seq peak summit coordinates were categorized according to their overlap with recruitment element on the X (*rex*, sites defined in (Albritton *et al*., 2017)), promoter [+ strand], promoter [- strand] (within 250 bp of a GRO-seq (Kruesi *et al*., 2013) or 500 bp of a Wormbase defined transcription start site (TSS)), active enhancer (overlapping a H3K27ac peak that is not a promoter), regulatory element (overlapping an ATAC-seq or DNase-seq peak and not promoter or active enhancer), and other, unknown categories. DPY-27 (DCC), H3K4me3, and H3K27ac wild type embryo ChIP-seq patterns are plotted across the DPY-27 ChIP-seq peak summits belonging to each category. **(E)** DCC, H3K4me3, H3K27ac ChIP-seq and ATAC-seq signals are plotted across X chromosome transcription start sites (defined by GRO-seq (Kruesi *et al*., 2013)); DCC signal coincides with the accessibility peak at promoters.

The current model for DCC binding to the X chromosomes involves two steps: recruitment and spreading (Csankovszki *et al*., 2004). Recruitment is mediated in a hierarchical manner, whereby the DCC enters the X at a small number of strong recruitment sites, which are fully distinguished from the autosomes by the presence of multiple copies of a 12-bp recruitment motif (McDonel *et al*., 2006; Ercan *et al*., 2007; Jans *et al*., 2009) within high occupancy transcription factor target (HOT) sites (Albritton *et al*., 2017). The stronger and weaker recruitment sites are thought to cooperate over long distances to robustly recruit the DCC to the X (Albritton *et al*., 2017). Unlike recruitment, spreading is an X-sequence independent process and can occur on DNA physically attached to the X (Ercan *et al*., 2009). An estimated 50-100 recruitment sites separated by 0.1-1 Mb distances support binding of the DCC across the ~17Mb X chromosome (Jans *et al*., 2009; Kranz *et al*., 2013; ALBRiTToN *et al*., 2017).

Global run on (GRO-seq) (Kruesi *et al*., 2013) and chromatin immunoprecipitation sequencing (ChIP-seq) (Kramer *et al*., 2015) analyses showed that the DCC is required to reduce RNA Pol II binding at X chromosomal promoters. DCC mediated repression appears to be chromosome-wide, with no large groups of genes escaping from dosage compensation (Kramer *et al*., 2015; Kramer *et al*., 2016). Previous work has highlighted multiple roles for the DCC in the regulation of X chromosome structure; DCC is required for the ~40% compaction of the X compared to autosomes (Lau *et al*., 2014), the regulation of subnuclear localization of the X chromosomes (Sharma *et al*., 2014; Snyder *et al*., 2016), and the regulation of topologically associating domains (TAD) on the X (Crane *et al*., 2015; Brejc *et al*., 2017). The DCC was also shown to increase and decrease the levels of H4K20me1 and H4K16ac, respectively, on the X chromosomes (Vielle *et al*., 2012; Wells *et al*., 2012). Reduction of H4K16ac on the X occurs downstream of H4K20me1 enrichment, and requires the deacetylase SIR-2.1 (Wells *et al*., 2012). H4K20me1 enrichment on the X is mediated by the H4K20me2 demethylase DPY-21, which physically interacts with the condensin core of the DCC (Brejc *et al*., 2017). How increased H4K20me1 and decreased H4K16ac mechanistically contribute to X chromosome repression is unclear (Kramer *et al*., 2015). in addition, previous studies did not address if the DCC regulates the level or the distribution of other histone modifications, such as H3K4me3 and H3K27ac that are tightly linked to transcription regulation.

To address this question, we analyzed the distribution of several histone modifications in wild type, DCC mutant and DCC depleted conditions, as well as in X-to-autosome fusion strains in which the DCC ectopically spreads into autosomal sequences (Ercan *et al*., 2009). in wild type embryos and L3 larval animals, DCC binding sites coincide with accessible putative gene regulatory elements marked by ATAC-seq (Daugherty *et al*., 2017). in DCC mutant (*dpy-21*, null) or depleted (*dpy-27*, RNAi) embryos, the levels of repressive histone modifications, including H3K9me3, remain unchanged while levels of active histone modifications, including H3K4me3 and H3K27ac, increase at X chromosomal promoters compared to autosomal ones. Further linking DCC binding to the regulation of active histone marks and gene expression, in X;V fusion chromosomes ectopic spreading of the DCC into autosomal sequence locally reduces both gene expression and H3K4me3. We also found that DCC depletion does not affect binding of PHA-4 transcription factor, the cohesin loader PQN-85 (Scc2 homolog), nor the putative H3K27acetylase CBP-1 as measured by ChIP-seq, thus ruling out a model in which DCC indiscriminately reduces binding of proteins to the X chromosomes. Taken together, our results are consistent with a model in which the DCC fine-tunes transcription across the X through targeting and modulating, directly or indirectly, the activity of gene regulatory elements by reducing the levels of specific activate histone modifications.

## RESULTS

### DCC is preferentially enriched at active gene regulatory elements on the X

To understand the DCC’s effect on histone modifications, we first compared the genomic distribution of the DCC to marks of active and repressed chromatin in embryos and L3 larval stages (Figure 1B). A combination of new and published ChIP-seq data, including those from modENCODE (Gerstein *et al*., 2010), and published accessibility data for ATAC-seq (Daugherty *et al*., 2017) and DNase-seq (Ho *et al*., 2017) were used. New data were produced from at least two biological replicates that correlate based on visual examination of genome browser tracks (Supplemental Figure 1). Summary and access information on all data sets is provided in Supplemental File 1.

Genome browser analysis of DCC binding revealed a correlation with active chromatin (Figure 1B) as previously noted (Ercan *et al*., 2007; Jans *et al*., 2009). We used data from more recent studies to refine the comparisons. DCC binding sites coincide with ATAC-seq peaks at promoters and enhancers containing RNA Pol II, H3K4me3 and H3K27ac (Figure 1B) (Daugherty *et al*., 2017). Conversely, marks of repressive chromatin, including H3K27me3 and H3K9me3 do not coincide with DCC binding (Figure 1B). Comparison of additional DCC subunits DPY-30 and SDC-3, histone modifications, and proteins including the transcription factor PHA-4, cohesin loader subunit PQN-85 (Scc2 homolog), putative H3K27 acetylase CBP-1 (p300 homolog), and the mediator subunit MDT-15 (Med15 homolog) support the conclusion that DCC binding coincides with gene regulatory sites (Supplemental Figure 2). To further refine which proteins the DCC best correlates with, we plotted the Spearman rank correlation of average ChIP-seq enrichment within 1 kb contiguous windows across the X chromosome (Figure 1C). DCC subunit DPY-27 binding correlates best with H3K4me3, RNA Pol II, MDT-15, and CBP-1.

### The complex pattern of DCC ChIP-seq profile suggests different modes of binding

The DCC has a complex pattern of binding as measured by ChIP-seq, including somewhat uniform baseline enrichment across the X, peaks of different heights at promoters, enhancers and within genes, and strong enrichment at the recruitment sites (Ercan *et al*., 2007; Albritton *et al*., 2017). To further categorize the sites of DCC enrichment, we focused on the top 50% of peaks sorted by their ChIP-seq score at the summit, mostly eliminating peak calls due to baseline DCC binding (Supplemental Figure 3). Next, we categorized the DCC peaks as those located at recruitment sites (Albritton *et al*., 2017), promoters (within 250 bp of a GRO-seq or 500 bp of a Wormbase defined transcription start site (TSS) (Kruesi *et al*., 2013)), active enhancers (overlapping a H3K27ac peak that is not a promoter), gene regulatory elements (overlapping an ATAC-seq or DNase-seq peak and not promoter or active enhancer), and unknown categories. We then plotted DCC, H3K4me3 and H3K27ac enrichment patterns across the DCC summits in each category (Figure 1D). This analysis revealed that a majority of DCC binding peaks occur at active promoters and enhancers, and that the strength of DCC binding correlates with the activity of the gene regulatory site, as measured by H3K4me3 and H3K27ac enrichment. DCC binding at promoters coincides with chromatin accessibility, and surrounding H3K27ac and H3K4me3 enrichment at the +1 nucleosome (Figure 1E). Approximately one third of DCC binding sites show little H3K4me3 and H3K27ac enrichment (Figure 1D), suggesting that DCC binding is not restricted to elements marked by these modifications.

### DCC binding at promoters partially correlates with their transcriptional activity

To understand the different modes of DCC binding, we next scrutinized the level of correlation between DCC binding and transcription, as shown by ChIP-chip analysis comparing DPY-27 and RNA Pol II in embryos and L4/young adults (Ercan *et al*., 2009). Similarly, DCC and RNA Pol II binding (as measured by ChIP-seq using 8WG16 antibody recognizing the unmodified C terminal of AMA-1 (large subunit)) show DCC enrichment changes at genes differentially bound by RNA Pol II in embryos and L3s (Figure 2A). Supporting the conclusion that DCC correlates with active transcription, DCC binding at promoters is higher at genes that are being transcribed compared to silent genes and genes whose mRNAs were maternally deposited in embryos (Figure 2B).

**Figure 2.**
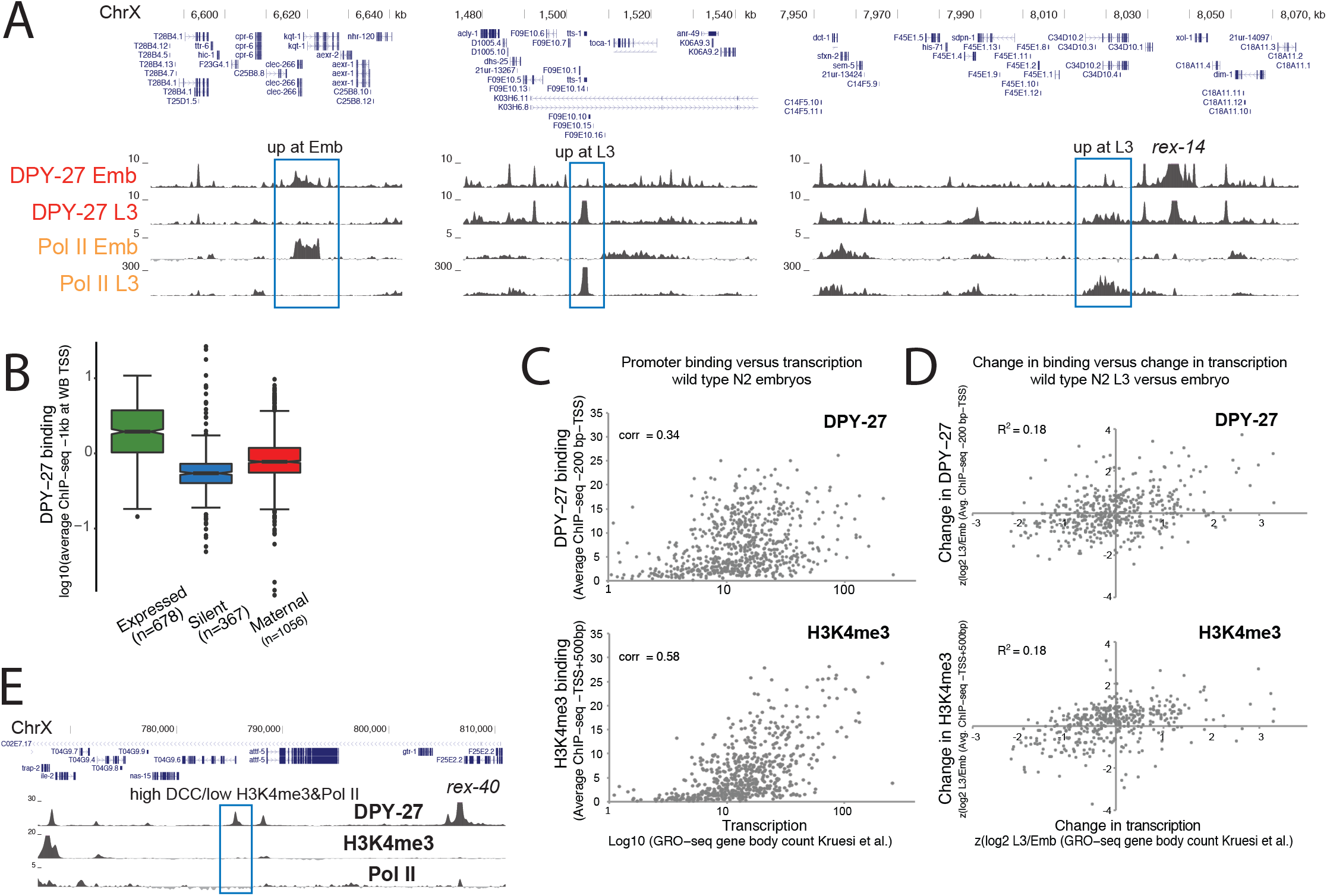
DCC enrichment at promoters partially correlates with transcriptional activity. **(A)** DPY-27 ChIP-seq binding across example X chromosomal regions with differential transcription as shown by RNA Pol II ChIP-seq in embryos versus L3s. **(B)** Average DPY-27 ChIP-seq score at 1kb windows centering around the X chromosomal Wormbase-defined TSS sites were plotted. Genes were categorized as expressed (N2 embryos FPKM >1 (Kramer *et al*., 2015) and detected in GRO-seq (Kruesi *et al*., 2013)), silent (FPKM = 0 and *not*, detected in GRO-seq), and maternally loaded (FPKM >1 and *not*, detected in GRO-seq). **(C)** Average DPY-27 and H3K4me3 ChIP-seq scores at proximal promoters (200 bp downstream to – TSS defined by (Kruesi *et al*., 2013)) were plotted on the y axis, and transcription level of genes (GRO-seq counts at corresponding gene bodies (Kruesi *et al*., 2013)) were plotted on the x axis. Spearman rank correlation coefficients are shown on the top left of each plot. **(D)** Changes in DPY-27 binding at promoters on the y axis (z score of log2 L3/embryo ratio of average ChIP-seq score within proximal promoters as in panel C) were compared to changes in transcription on the x axis (z score of log2 L3/embryo of transcription level as in panel C) in L3 versus embryos. Change in DPY-27 and H3K4me3 partially correlates with the change in transcription at individual promoters. **(E)** UCSC browser view of DPY-27, H3K4me3, RNA Pol II ChIP-seq signal across a 40 kb region containing a recruitment site. The DCC binding peak highlighted with a blue rectangle shows low Pol II and H3K4me3, suggesting that DCC enrichment and transcriptional activity at promoters can be uncoupled.

However, the level of positive correlation between DCC and transcription (as measured by GRO-seq (Kruesi *et al*., 2013), Spearman rank correlation of 0.34) is less than that observed between H3K4me3 and transcription (0.58), suggesting that the link between DCC and RNA Pol II binding is weaker than that of H4K3me3 (Figure 2C). To evaluate how DCC and H3K4me3 are tuned to transcription at individual promoters, we plotted the change in DCC or H3K4me3 levels versus change in transcription between embryos and L3s (Figure 2D). While there is a slight positive correlation, both DCC and H3K4me3 do not perfectly follow transcription changes at individual promoters. Furthermore, at and near recruitment sites, we found sites with high DCC and low H3K4me3 and RNA Pol II (Figure 2E). These results suggest that while DCC binding generally correlates with transcriptional activity, the two are not strictly coupled.

### DCC reduces the levels of active histone modifications on the X

To determine DCC’s effect on gene regulatory elements, we analyzed several histone modifications associated with active and repressed chromatin upon DCC knockdown (*dpy-27*, RNAi) and mutation (*dpy-21(e428) V)*, in embryos. Since DCC represses transcription by approximately 2-fold, we expected and observed subtle changes. To quantify such subtle changes, we used the autosomes as an internal control for ChIP efficiency and calculated the standardized ratio of ChIP enrichment in mutant versus wild type. This approach detected previously described X-specific changes upon DCC mutation and knockdown, including for H4K20me1, H4K16ac (Vielle *et al*., 2012; Wells *et al*., 2012) (Figure 3A) and RNA Pol II (Pferdehirt *et al*., 2011; Kramer *et al*., 2015) (Figure 3B). The levels of histone H3 and negative control IgG at promoters do not show a significant change (Figure 3C), ruling out a nonspecific effect on nucleosome occupancy. We then applied the same analyses to additional histone modifications and found a DCC-dependent reduction in the X chromosomal levels of H3K4me3 (Figure 3D), H3K27ac (Figure 3E), and H4pan-ac (K5,8,12,16), but not H3ac (Figure 3F), suggesting that DCC activity correlates with a reduction in specific active histone modifications at X chromosomal promoters.

**Figure 3.**
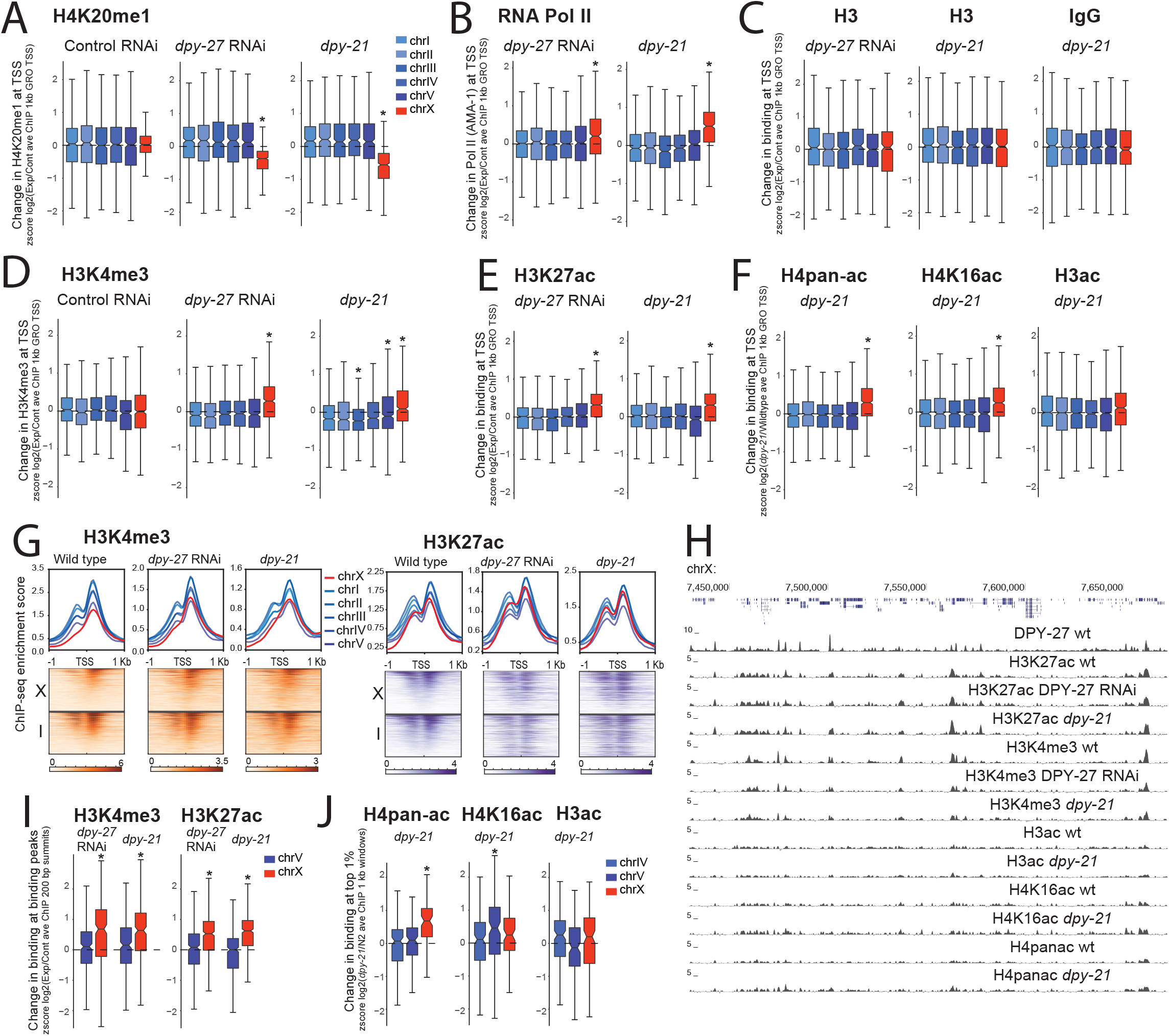
DCC is required for reduction of active histone modifications on the X. Changes in levels of histone modifications upon DCC defect in embryos are plotted. Average ChIP enrichment within 1 kb windows centered at the GRO-seq defined transcription start sites (Kruesi *et al*., 2013) was calculated in wild type (N2), DCC mutant (*dpy-21*,) and DCC depleted (*dpy-27 RNAi*,) embryos. Change in the level of each histone modification was measured by standardizing (z score) log2 ratio of experimental to control ChIP-seq scores. Values from each chromosome were tested against all the other autosomes using a two-tailed Student’s t-test and resulting p-values that were less than ≤ 0.001 were marked with an asterisk. This analysis captured the expected changes in H4K20me1 **(A)** and RNA Pol II **(B)** at X chromosomal promoters upon DCC defect. **(C)** Neither H3 nor IgG negative control ChIP-seq data showed a comparable difference in the *dpy-21*, mutant, and H3 in *dpy-27*, RNAi suggesting that nucleosome levels are not significantly affected. **(D-F)** Same analysis of different histone modifications associated with active transcription. Note that H4 panac antibody also recognizes H4K16ac, thus changes may be due to this modification. **(G)** ChIP-seq enrichment for H3K4me3 and H3K27ac in wild type, *dpy-21*, mutant and *dpy-27*, RNAi knock down embryos was plotted across the GRO-seq defined transcription start sites (Kruesi *et al*., 2013) on the X and chromosome I. The level of enrichment is ordered in a descending manner using maximum coverage in wild type. Mutant data was plotted in the same order. **(H)** Genome browser view of ChIP-seq profiles in wild type, mutant and knockdown embryos over a 250kb representative region of the X chromosome. The pattern of enrichment in wild type and mutants is largely similar. **(I)** As in (A), but change in binding at the wild type peak summit, rather than TSS. Standardized (z score) log2 ratio of mutant/wild type ChIP-seq score within a 200 bp window centering at the summit of peaks in wild type embryos. **(J)** Similar analysis as in (I), but change in top 1% of 1kb windows ordered by average ChIP-seq in wild type embryos.

DCC depletion caused an increase in active histone modifications at their canonical locations rather than changing their distribution (Figure 3H). H3K4me3 and H3K27ac enrichment across the transcription start sites in wild type, *dpy-21*, mutant and *dpy-27*, RNAi conditions show small differences, but generally, sites with high enrichment in the wild type are still highly enriched in the mutant conditions (Figure 3G). Furthermore, Spearman rank correlation values between wild type and mutant H3K4me3 and H3K27ac enrichment within 1 kb contiguous windows were similar on the autosomes and the X (H3K4me3 N2-CB428 on X: 0.59 on autosomes:0.57-0.65; H3K4me3 control-*dpy-27*, RNAi on X:0.55 on autosomes:0.52-0.65; H3K27ac N2-CB428 on X:0.79 on autosomes:0.79-0.85; H3K27ac N2-*dpy-27*, RNAi on X:0.85 on autosomes:0.84-0.89), indicating a lack of X-specific change in the distribution of histone modifications upon DCC defect. Collectively, these results suggest that DCC depletion did not create or eliminate new sites of enrichment on the X but increased the level of H3K4me3 and H3K27ac at their canonical locations.

Since the distribution of modifications remains similar in the DCC knockdown embryos, we analyzed the level of change at their canonical sites (ChIP-seq peaks in wild type). The levels of H3K4me3 and H3K27ac within 200 bp of their canonical binding summits increase specifically on the X upon DCC defect (Figure 3I). Peak calling on lower and broader ChIP-seq patterns observed for H4pan-ac, H4K16ac, and H3ac was difficult, therefore to analyze their binding, we took the top 1% of 1 kb windows based on wild type ChIP-enrichment. in the *dpy-21*, mutant, the level of H4pan-ac increases specifically on the X, but H4K16ac and H3ac do not (Figure 3J). Greater variability in the *dpy-21*, mutant compared to *dpy-27*, RNAi may be due to additional *dpy-21*, activity outside dosage compensation (Kramer *et al*., 2015; Brejc *et al*., 2017). For H4K16ac, an X-specific effect at the TSS (Figure 3F), but not at top 1% H4K16ac sites (Figure J) suggests spatial specificity for DCC–mediated reduction of H4K16ac, possibly by SIR-2.1, which was shown to be required for H4K16 deacetylation on the X (Wells *et al*., 2012).

Since DCC reduces RNA Pol II binding and active histone modifications on the X (Kruesi *et al*., 2013; Kramer *et al*., 2015), we asked if the DCC-dependent decrease in Pol II binding correlates with decrease in histone modifications at individual promoters. RNA Pol II and DCC binding positively correlate with the levels of active histone modifications at promoters in both wild type and *dpy-21*, mutant embryos (Supplemental Figure 4A). At individual promoters on the X, change in RNA Pol II binding did not correlate as well with the change in active histone modifications (Supplemental Figure 4B). Nevertheless, RNA Pol II and H3K4me3 ChIP-seq ratios in *dpy-21*, mutant versus wild type positively correlated (0.21 for AMA-1 antibody). Interestingly, DPY-27 and H3K27ac in *dpy-21*, versus wild type negatively correlated specifically on the X (Supplemental Figure 4B, spearman rank correlation of −0.32 on X and 0.04 on autosomes), supporting the idea that the DCC is linked to a reduction in H3K27ac on the X chromosomes.

### DCC does not affect repressive histone modifications

To test if the DCC affects histone modifications indiscriminately, we performed ChIP-seq analysis of H3K4me1, H3K27me1, H3K27me2 and H3K9me3 (Figure 4A). H3K4me1, H3K27me1 and H3K27me2 did not yield strong signals, precluding clear conclusions (Supplemental Figure 1). Nevertheless, changes in these modifications were neither X-specific nor consistent between *dpy-27*, RNAi and *dpy-21*, mutant (Supplemental Figure 5A). We observed no difference in H3K9me3 distribution on the X chromosomes between control and *dpy-27*, RNAi treated embryos (Figure 4A). H3K9me3 levels showed higher variability in control and *dpy-27*, RNAi conditions, but the difference is not restricted to the X chromosome suggesting that RNAi treatment affects H3K9me3 across the genome (Figure 4B). We also considered the opposite, and tested if H3K9me3 affects DCC localization by using a strain in which H3K9 methylation is eliminated (Towbin *et al*., 2012). in the absence of H3K9me3, RNA Pol II binding pattern is similar to that of wild type (Figure 4C), consistent with the lack of overt effect on growth in laboratory conditions (Towbin *et al*., 2012). The reason for a general reduction in ChIP scores in the mutant is unclear. Regardless, RNA Pol II binding is not specifically different on the X (Figure 4D), consistent with there being no strong link between H3K9me3 and the DCC. Similarly, SDC-3 (DCC subunit required for DPY-27 recruitment to the X), and CAPG-1 (HEAT domain subunit of condensin DC) binding profiles are similar between the wild type and mutant (Figure 4C). Furthermore, CAPG-1 peaks in the H3K9me3 mutant largely overlap with those of the wild type (Figure 4E), and genomic sites with high H3K9me3 enrichment do not coincide with new SDC-3 sites (Figure 4F). Collectively, these results suggest that DCC does not regulate and is not regulated by the heterochromatin mark H3K9me3.

**Figure 4.**
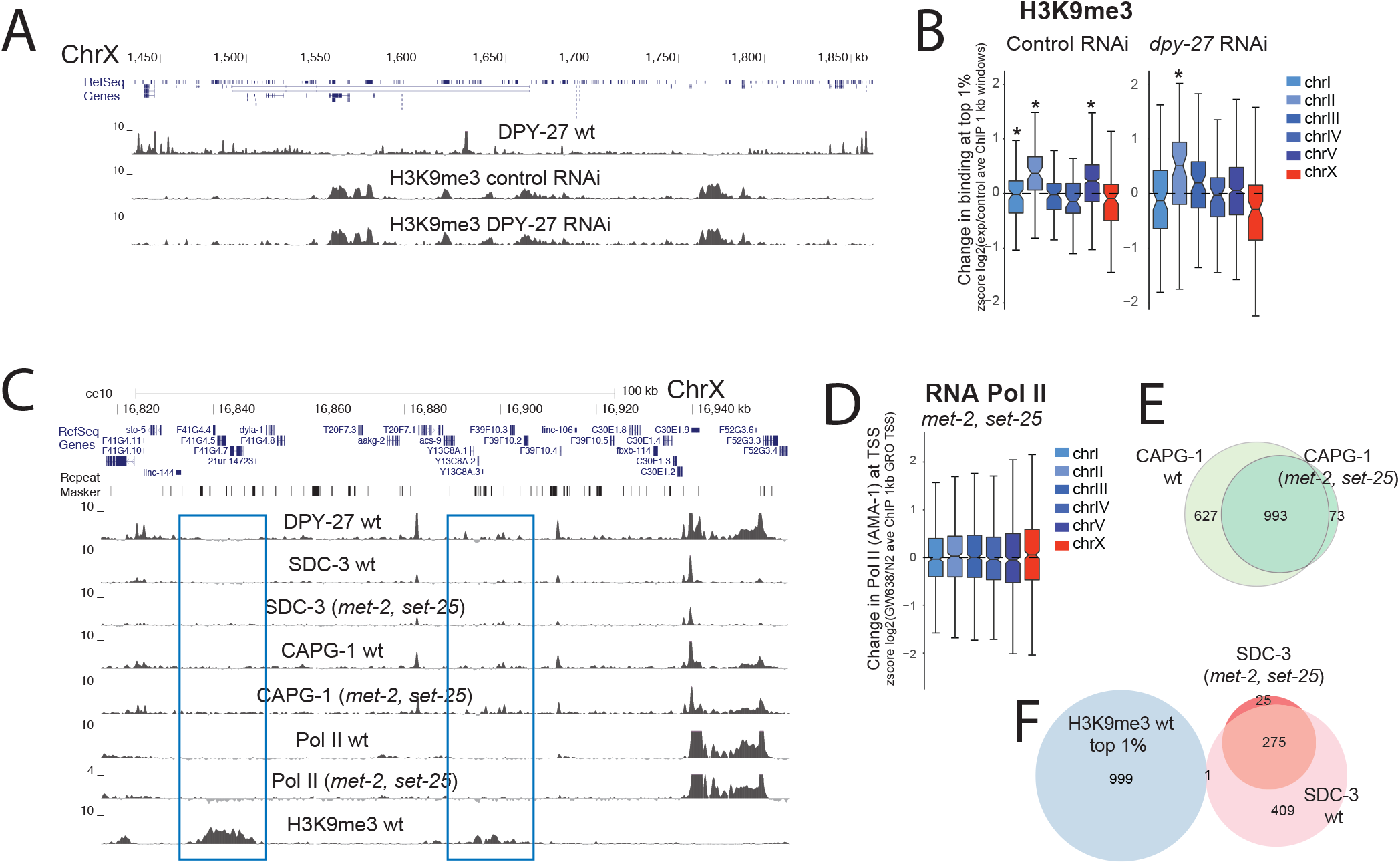
DCC does not affect the levels of repressive histone marks. **(A)** ChIP-seq profile of H3K9me3 in control and DPY27 RNAi embryos along a representative region of chr X exemplifies no significant change in H3K9me3 upon DCC knockdown. **(B)** Distribution of standardized (z score) log2 *dpy-27*, RNAi/N2 ratios of H3K9me3 ChIP-seq average at the top 1% most enriched 1kb windows. Values from each chromosome were tested against all the other autosomes using a two-tailed Student’s t-test and resulting p-values that were less than ≤ 0.001 were marked with an asterisk. **(C)** ChIP-seq profiles of DPY-27, DCC subunits (SDC-3, CAPG-1), Pol II, and H3K9me3 in wild type and the H3K9me3 null mutant (GW638, *met-2, set-25*,) across a representative region of the X. DCC and Pol II binding profiles remained similar in the *met-2, set-25*, mutant, including in regions enriched in H3K9me3 in wild type (blue rectangle). **(D)** Pol II binding in *met-2, set-25*, H3K9me3 null mutant (GW638) compared to N2 wild type showed no specific effect on X chromosome expression. Distribution of standardized (z score) log2 mutant/N2 ratio of RNA Pol II ChIP-seq within 1 kb windows centered at the GRO-seq defined transcription start sites (Kruesi *et al*., 2013). **(E)** ChIP-seq peak overlap of DCC subunit CAPG-1 between wild type and H3K9me3 null mutant. **(F)** ChIP-seq peak overlap between SDC-3 and top 1% H3K9me3 enriched 1kb windows.

### DCC spreading into autosomal loci in X;A fusion chromosomes represses gene expression

To determine if DCC spreading reduces gene expression and histone modification levels locally, we analyzed strains containing X-to-autosome fusion (X;A) chromosomes. Previous work showed that the DCC spreads into the autosomal regions of X;V, X:II, and X:I fusion chromosomes, and that spreading is linear and reduces with distance from the X (Ercan *et al*., 2009). Ectopic DCC binding leads to increased H4K20me1 in the autosomal region of spreading (Vielle *et al*., 2012). An earlier microarray analysis in embryos did not detect a significant difference in gene expression in the fusion strains (Ercan *et al*., 2009). Subsequent experiments suggested that dosage compensation starts in embryogenesis but is not complete until larval stages (Kramer *et al*., 2015), therefore we repeated the experiment using mRNA-seq in larvae.

DPY-27 ChIP-seq analysis verified that the DCC binds to the autosomal region of spreading in the X;V fusion chromosomes in larvae (Figure 5A). To test if gene expression specifically changed in the region of spreading, first we took three 0.5 Mb windows at the middle, left- and right-most end of each chromosome, and plotted the ratio of mRNA-seq levels for genes within each window. Average gene expression is significantly and specifically reduced at the side of X fusion, which is the right most end of chromosome V in the X;V strain, and the left-most end of chromosome II in the X;II strain (Figure 5B). The level of repression reduces with distance from the fusion site (Figure 5C), and is proportional to the level of spreading demonstrated for each fusion (Ercan *et al*., 2009). These results indicate that DCC spreading into the autosomal regions of the X;V and X;II fusion strains results in repression.

**Figure 5.**
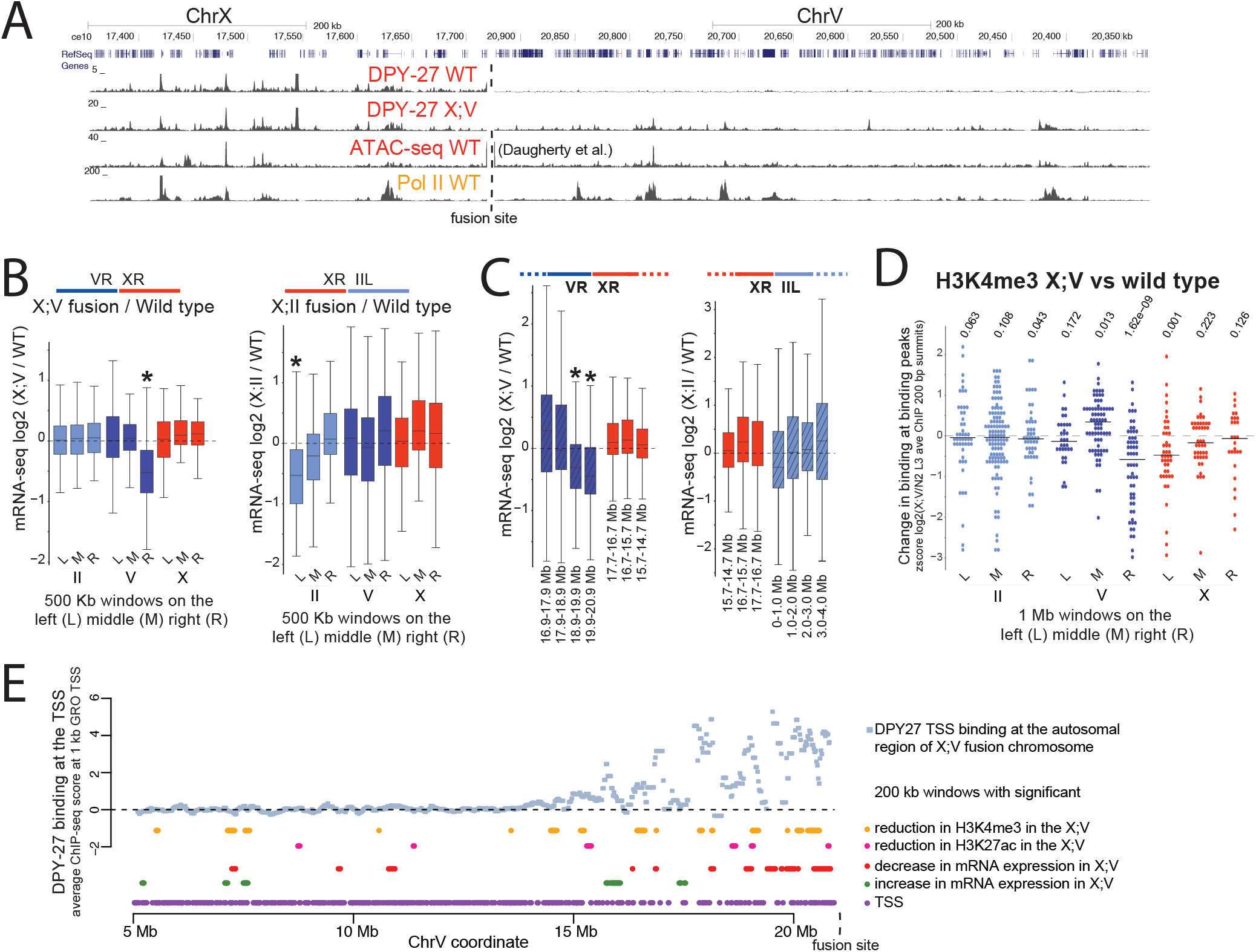
DCC spreading into X;A fusion chromosomes reduces gene expression. **(A)** DPY-27 (DCC) ChIP-seq profile in the wild type and X;V fusion chromosome containing strains in L3 larvae. The spreading profile of the DCC in the autosomal region of the fusion chromosome is similar to that on the X, as indicated by Pol II ChIP-seq and ATAC-seq signal in wild type. **(B)** mRNA-seq analyses in wild type and strains containing X;V and X;II fusion chromosomes. DESeq log2 expression ratios were calculated and plotted for genes located within middle, left and right-most 500kb windows of chromosomes II, V, and X. * p-value ≤ 0.001 (two-tailed Fisher test for each window against the rest of the windows across the genome). The schematics above the boxplots show which chromosome arms are fused. For X;II, the right end of X was fused to the left end of chr II, and for X;V the right end of X was fused to the right end of V (Lowden *et al*., 2008). **(C)** Similar to panel B, but expression ratios were plotted for genes within 1 Mb windows stepping out from the fusion site. The amount of repression decreases as a function of distance from the fusion border, following the pattern of DCC spreading (Ercan *et al*., 2009). **(D)** Change in H3K4me3 levels in X:V fusion chromosome compared to wild type at the middle, left and right-most 1Mb windows of chromosomes II, V, and X. Standardized (z score) log2 X;V/wild type ratios of ChIP-seq score within 200bp H3K4me3 peak summits were plotted. H3K4me3 slightly but significantly decreases in the DCC spreading region (two-tailed Student’s t-Test comparing ratios of each 1 Mb window against the rest across the genome). **(E)** Average DPY-27 ChIP-seq scores for 1 kb windows centering at GRO-seq defined transcription start sites (Kruesi *et al*., 2013). Changes in expression and histone modifications were calculated by a moving average analysis using a 200 kb window with a 20 kb step size. For each 200 kb window, ChIP-seq and mRNA-seq ratios in X;V/wt were compared to the rest of the chromosome and a p-value statistic was generated through t-test. In this analysis rather than asking if there is a significant change for each gene (as in DEseq), we ask whether the values in each window are higher or lower than the values observed for the rest of the windows along the chromosome. Windows with a p-value ≤ 0.01 are clustered towards the region of spreading.

### DCC spreading into autosomal loci leads to H3K4me3 reduction

In the X;V fusion strain, DCC spreads further (Ercan *et al*., 2009) and causes stronger repression (Figure 5C). Thus, we assayed the change in H3K4me3 and H3K27ac levels in the X;V fusion chromosomes. Since the expected change is small, we focused the analysis on where the signal is highest by taking a standardized ratio of ChIP-seq enrichment in X;V versus wild type at 200 bp around canonical peak summits. Despite high variability, there is a significant reduction in average H3K4me3 levels within the spreading domain of the X;V fusion chromosomes compared to wild type (Figure 5D). For H3K27ac, there is higher variability across chromosomes, yet a slight reduction within the autosomal region of spreading is also observed (Supplemental Figure 6A).

To be able to analyze DCC spreading with respect to the subtle changes in gene expression and histone modification levels, we used a sliding window analysis with 200 kb windows and 20 kb steps. For each window, we performed a student’s t-test asking whether the ChIP-seq or mRNA-seq ratio within each window is significantly increased or decreased compared to the rest of the windows across chrV. Windows with p values less than 0.01 were plotted under the DPY-27 ChIP-seq enrichment in the X;V fusion chromosome (Figure 5E). Although noisy, the level of gene expression, H3K4me3 and H3K27ac were slightly reduced in windows close to the fused end of chrV (Figure 5E). Lack of a similar pattern on other chromosomes (Supplemental Figure 6B) supports the conclusion that DCC spreading into the autosomal region of the fused chromosome reduces active histone modifications and represses transcription.

### DCC depletion does not significantly alter binding of PHA-4, CBP-1 and PQN-85

To test if the DCC represses transcription by reducing binding of all proteins to the X chromosomes, we performed ChIP-seq analysis of the transcription factor PHA-4, the putative H3K27 acetylase CBP-1 (p300 homolog), and the cohesin loader subunit PQN-85 (Scc2 homolog) (Figure 6A). We found no X-specific difference in the binding of these proteins as measured by ChIP-seq upon *dpy-27*, RNAi knockdown (Figure 6B), suggesting that the DCC does not indiscriminately displace proteins from the X.

**Figure 6.**
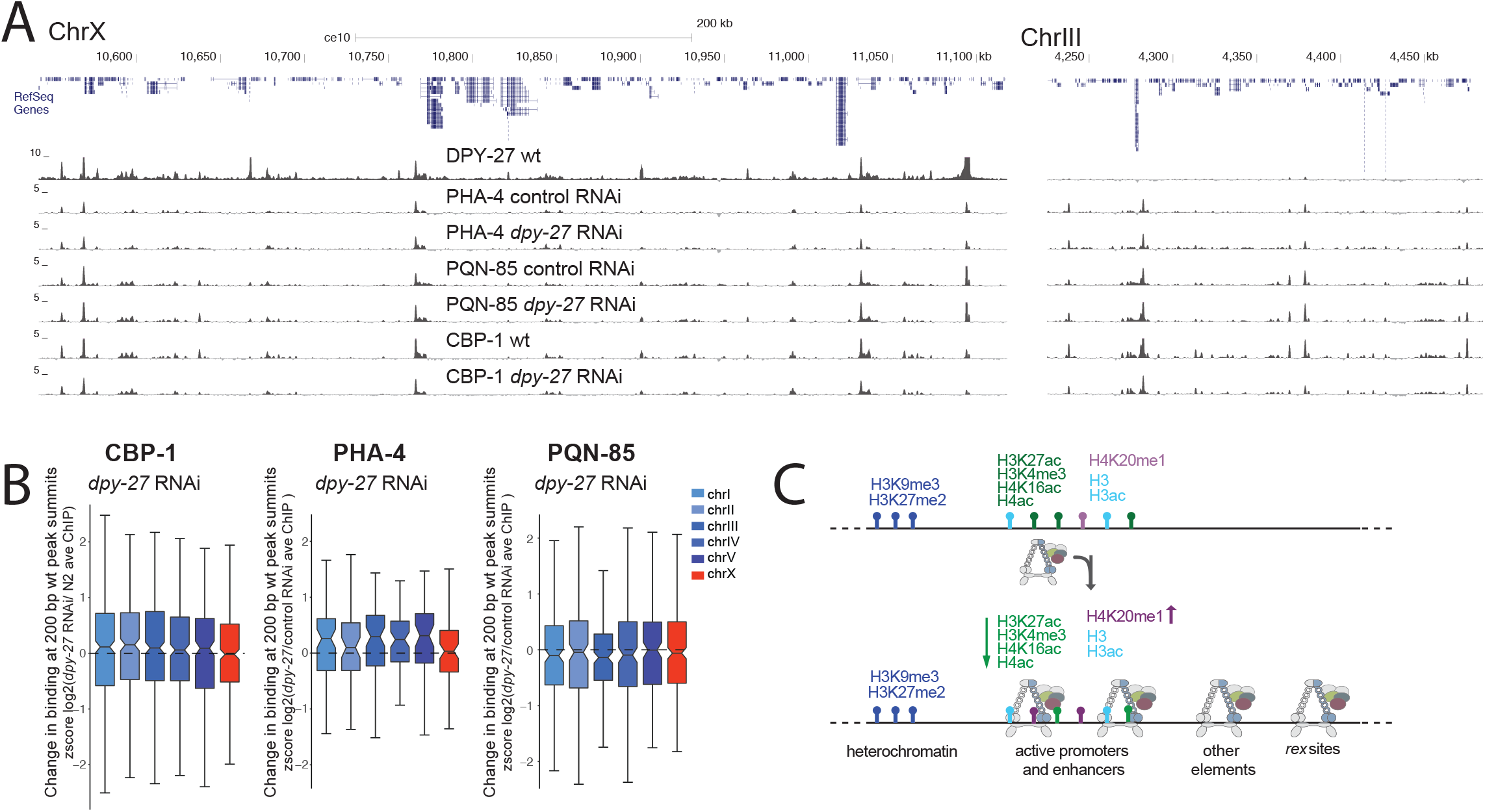
DCC knockdown does not indiscriminately reduce protein binding as measured by ChIP-seq. **(A)** ChIP-seq profiles of DPY-27 (condensin DC subunit), PHA-4 (FOXA transcription factor), PQN-85 (*S. cerevisiae*, Scc2p homolog), and CBP-1 (putative H3K27 acetyltransferase), in representative regions on chr X and III. **(B)** Analysis as in figure 3I, plotting change in protein binding across 200 bp wild type peak summits. CBP-1, PQN-85, and PHA-4 levels on the X chromosomes did not change significantly upon DCC knockdown. **(C)** Summary of DCC binding and regulation of histone modifications on the X chromosomes. DCC binding sites coincide with gene regulatory elements marked by accessible chromatin on the X. The majority of these elements also contain histone modifications associated with active transcription. The remaining include recruitment elements and sites that do not contain the analyzed histone modifications. DCC activity correlates with X-specific changes in the level of specific histone modifications (denoted by up and down arrows).

## DISCUSSION

Our analysis of DCC distribution with respect to various chromatin marks reflects multiple modes of binding, including a baseline distribution, strong enrichment at the recruitment sites, and peaks of DCC enrichment at gene regulatory elements, partially correlating with transcription. our results suggest that the DCC is required for reducing the level of histone modifications that are associated with active transcription, including H3K4me3 and H3K27ac, but does not regulate nor is regulated by the heterochromatin-associated histone modification H3K9me3. GRO-seq and ChIP-seq analysis of transcription in DCC mutants showed that the DCC reduces RNA Pol II binding to X chromosomal promoters (Pferdehirt *et al*., 2011; Kruesi *et al*., 2013; Kramer *et al*., 2015). Collectively, these results suggest a model in which DCC binding is directly or indirectly linked to a reduction in the activity of X chromosomal gene regulatory elements (Figure 6C).

The first question this model raises is how the DCC targets active gene regulatory elements? Accumulation at active promoters and enhancers is a conserved feature of condensins in *C. elegans*,, *D. melanogaster*,, chicken, mouse and human cells (Jeppsson *et al*., 2014). Condensins bind chromosomes by entrapping and/or encircling DNA through multiple interactions mediated by its ring structure (Cuylen *et al*., 2013; Kschonsak *et al*., 2017). one mechanism by which condensins may target gene regulatory elements is through binding to accessible DNA *in vivo*,, which tends to coincide with active promoters and enhancers. Another possibility is through specific recruitment by transcription factors. in yeast and mammals, condensins are recruited to tRNA gene promoters and extra TFIIIC sites by interacting with TFIIIC (D’Ambrosio *et al*., 2008; Haeusler *et al*., 2008; Iwasaki *et al*., 2010; Kranz *et al*., 2013; Van Bortle *et al*., 2014; Yuen *et al*., 2017), TBP (Iwasaki *et al*., 2015) and sequence specific transcription factors (Kim *et al*., 2016). in *C. elegans*,, the strong DCC recruitment elements are HoT sites that are bound by multiple transcription factors (Albritton *et al*., 2017). Binding to accessible DNA and recruitment by specific transcription factors are not mutually exclusive mechanisms (Robellet *et al*., 2017). Indeed, condensin DC binding through both DNA accessibility and specific recruiter proteins may result in the complicated pattern of DCC distribution that we observe *in vivo*.,

The second question that our model raises is how specific histone modifications are regulated by the DCC? Our work suggests that the DCC does not indiscriminately reduce binding of proteins to the X. DCC may recruit specific histone deacetylases, e.g. *sir-2.1*,, which is required to reduce H4K16ac on the X (Wells *et al*., 2012). The observation that the DCC reduces H3K27ac but not CBP-1 binding suggests that similar to H4K16ac, H3K27ac reduction may also depend on a deacetylase. It is also possible that the DCC regulates binding of specific histone modifying complexes. For instance, physical interaction of the DCC with a subunit of a chromatin-modifying complex may serve as a barrier. Supporting this idea, DPY-30, an essential subunit of the MLL/COMPASS complex physically interacts with the DCC (Pferdehirt *et al*., 2011). Intriguingly, a recent proteomic analysis found that mitotic chromosomes disproportionately lose chromatin-modifying complexes associated with euchromatin and not heterochromatin (Ginno *et al*., 2018). Furthermore, the level of displacement differed for different histone acetylases (Ginno *et al*., 2018). it is possible that reduced acetylation is connected to mitotic transcriptional repression, which is thought to be important for chromosome segregation (Sutani *et al*., 2015). Therefore, DCC mediated transcriptional repression may have evolved from a conserved condensin role in regulating specific chromatin modifying complexes in the formation of mitotic chromosomes.

The third question is how do histone modifications regulate RNA Pol II binding to X chromosomal promoters? H3K4me3 and H3K27ac are particularly instructive in models that predict gene expression from histone modifications (Gerstein *et al*., 2010; Karlic *et al*., 2010; Zhang and Zhang 2011). We also observed a strong correlation between RNA Pol II, H3K4me3 and H3K27ac. At individual promoters, the differential binding of RNA Pol II and histone modifications upon DCC defect was less correlated, perhaps due to insufficient sensitivity of the ChIP-seq assay in *C. elegans*, embryos and/or a complex quantitative relationship between Pol II recruitment and histone modifications at a given promoter (Perez-Lluch *et al*., 2015). While it remains unclear if and how much H3K4me3 activates transcription directly (Howe *et al*., 2017), it has been shown that H3K4me3 interacts with specific transcriptional activators (Howe *et al*., 2017), and ectopic recruitment of H3K4me3 activates and maintains transcription (Cano-Rodriguez *et al*., 2016). H3K27ac is also associated with transcription activation, presumably by controlling transcription factor binding and RNA Pol II release from promoters (Stasevich *et al*., 2014). in *C. elegans*,, a small proportion of genes show promoter pausing (Maxwell *et al*., 2014), thus H3K27ac may regulate dynamics of activator binding upstream of RNA Pol II recruitment to promoters. Recent work using histone mutants in *D. melanogaster*, suggest that H3K27ac is not required for transcription (McKay *et al*., 2015; Leatham-Jensen *et al*., 2019); thus future work is required to determine how instructive H3K27ac is for transcriptional activation in different systems.

Evolution of diverse dosage compensation strategies reveals how different transcriptional regulatory mechanisms can be co-opted to regulate large domains within the genome. DCC belongs to the deeply conserved SMC family of complexes that are involved in genome organization and gene regulation across species (Hirano 2006; Dowen and Young 2014; Rowley and Corces 2018). Here, we show that the DCC targets gene regulatory elements and its binding correlates with changes in the level of active histone modifications rather than their distribution, suggesting that *C. elegans*, dosage compensation evolved to control transcriptional output without interfering with the underlying transcriptional program. A similar condensin-mediated tuning of histone modifications on mitotic chromosomes may be important for proper inheritance of transcriptional programs after cell division. Whether the changes in histone modifications are a cause or consequence of transcriptional repression is an important open question. Understanding how the DCC directly or indirectly modulates histone modifications and transcriptional activity of gene regulatory elements will help reveal mechanisms by which condensin-mediated organization of mitotic chromosomes affects gene regulation across cell division.

## MATERIALS AND METHODS

### Worm strains and growth

Mixed developmental stage embryos (wild type N2) were isolated from gravid adults by bleaching. Mutant strains used in this study were CB428 (*dpy-21(e428) V*,), oP37 (*wgIs37 [pha-4::TY1::EGFP::3xFLAG + unc-119(+)]*,), YPT41 (*X;II*,) and YPT47 (a.k.a. 15eh#1, X;V) (Lowden *et al*., 2008), and GW638 (*met-2(n4256) set-25(n5021) III*,) (Towbin *et al*., 2012). For ChIP samples, embryos or larvae were incubated in 2% formaldehyde for 30 minutes. Synchronized L3 worms were isolated by growing starved L1s for 24 hours at 22°C. L1-L3 worms were isolated from asynchronous plates by passing larvae with a 20-micron filter, where the embryos and larvae with expanded germline are not capable of flowing through. Large scale RNAi knockdown for ChIP and RNA-seq analyses was performed as described previously (Kranz *et al*., 2013). Briefly, bacteria with RNAi inducing plasmids were grown in liquid, and concentrated 130-fold to seed 6×10 cm plates. Synchronized N2 L1s were plated on RNAi plates and grown at 20°C for four days to obtain gravid adults. Knockdown was verified by western blot analysis of DPY-27 compared to control (vector only) RNAi. In previous work, we found that knockdown of DPY-27 in embryos isolated from mothers that were fed RNAi bacteria was more efficient than knockdown in L3s isolated after feeding L1s for one day (Kramer *et al*., 2015), thus RNAi experiments were performed in embryos.

### Antibodies and chromatin immunoprecipitation

Experiments were from at least two biological replicates with matching input samples as reference (Supplemental File 1). ChIP-seq (Kranz *et al*., 2013) and mRNA-seq (Albritton *et al*., 2014) experiments were performed as previously described. Information on antibodies used in this study is given in Supplemental File 1. Two new antibodies were used. MDT-15 antibody was validated by western blot analysis upon RNAi knockdown and immunoprecipitation (Supplemental Figure 7). CBP-1 antibody did not show a measurable signal on western blot hybridization and immunofluorescence assays but showed the expected ChIP-seq pattern overlapping with H3K27ac (Supplemental Figure 1), and immunoprecipitated CBP-1 specifically as analyzed by mass spectrometry. Briefly, whole embryos extract was prepared by douncing and sonicating embryos (5 min, 30 sec on 30 sec off in Bioruptor) in lysis buffer (40 mM HEPES, pH 7.5, 10% glycerol, 150 mM NaCl, 1 mM EDTA, 0.5% NP-40) complemented with protease inhibitors. After spinning insoluble material at 17,000 g for 15 min, 2mg of protein were incubated overnight with 5μg of rabbit polyclonal CBP-1 antibody and IgG as negative control, collected on protein A sepharose beads, washed five times using IP buffer (50 mM HEPES-KOH, pH 7.6, 1 mM EDTA, 150 mM NaCl), and subjected to trypsin digestion and mass spectrometry by the NYU Medical School Proteomics Facility on an Orbitrap Fusion Lumos. MS/MS spectra were searched against a Uniprot *C. Elegans*, database using Proteome Discoverer 1.4 (Supplemental File 1).

### ChIP-seq data processing

Single-end sequencing was performed by Illumina Genome Analyzer IIx, HiSeq-2000, HiSeq-2500, HiSeq-4000 or NextSeq 500. The raw and processed data are provided at Gene Expression Omnibus database (GEO, http://www.ncbi.nlm.nih.gov/geo) under accession number GSE122639. ChIP data processing and peak finding was performed as described previously (Kranz *et al*., 2013). Briefly, 50-75 bp single-end reads were aligned to the *C. elegans*, genome version WS220 using bowtie version 1.2.0 (Langmead *et al*., 2009), allowing two mismatches in the seed, returning the best alignment, and restricting multiple alignments to four sites in the genome. Mapped reads from ChIP and input were used to call peaks and obtain read coverage per base using MACS version 1.4.3 (Zhang *et al*., 2008) with default parameters. ChIP scores per base were obtained by normalizing to the median coverage and subtracting the input coverage. To obtain summits for binding profiles that are a combination of focused and broad patterns, large peaks were split using PeakSplitter version 1.0 (Salmon-Divon *et al*., 2010), with a minimum height cut-off of 4 and a separation float of 0.86. The replicate, number of reads, and access information for the data sets are provided in Supplemental File 1.

### ChIP-seq data analysis

Data were visualized using UCSC genome browser, ce10 (http://genome.ucsc.edu/). Heatmaps of ChIP enrichment across WS220 TSS and GRO-seq defined TSS sites (Kruesi *et al*., 2013) were produced using Deeptools (Ramirez *et al*., 2014) with default parameters in Galaxy (doi:10.1093/nar/gkw343). Change in ChIP binding scores across TSS and peak summits were calculated by standardizing average ChIP scores within a 1 kb window centering at the TSS or 200 bp window centered at the summit through calculating = (log2(mut/wt)-mean(log2(mut/wt)) / stdev (log2(mut/wt)). Box plots were produced in R using ggplot2 (http://ggplot2.org). The whiskers extend from the hinge to the largest value no further than +/-1.5 IQR (distance between the first and third quartiles) from the hinge. Outliers are not plotted. The notch shows the 95% confidence interval of the median (median +/-1.58*IQR/sqrt(n)). Data analysis scripts are available at the Ercan lab github: https://github.com/ercanlab/street_et_al_2019/.

### mRNA-seq data processing and analysis

Single-end sequencing was performed by Illumina HiSeq-2000. mRNA-seq data processing was performed as described previously (Albritton *et al*., 2017). Briefly, 50 bp single-end reads were aligned to the *C. elegans*, genome version WS220 using Tophat version 2.1.1 (Trapnell *et al*., 2012), using default parameters. Count data was calculated using HTSeq version 0.6.1 (Anders *et al*., 2015) and normalized using the R package DESeq2 (Anders and Huber 2010). The resulting mRNA levels and expression ratios are provided in Supplemental File1.

## ACKNOWLEDGEMENTS

We thank Susan Gasser for providing the GW638 strain, Dominic Balcon and Jacob Carmichael for help with growing worms and NYU-CGSB, UNC, and MDC Berlin High Throughput Sequencing Facilities for sequencing. We thank Gyorgyi Csankovszki for providing H4K16 antibody. Research reported in this publication was supported by NIGMS of the National Institutes of Health under award number R01GM107293. Some strains were provided by the CGC, which is funded by the NIH Office of Research Infrastructure Programs (P40 OD010440).

---

**Supplemental Figure 1.**
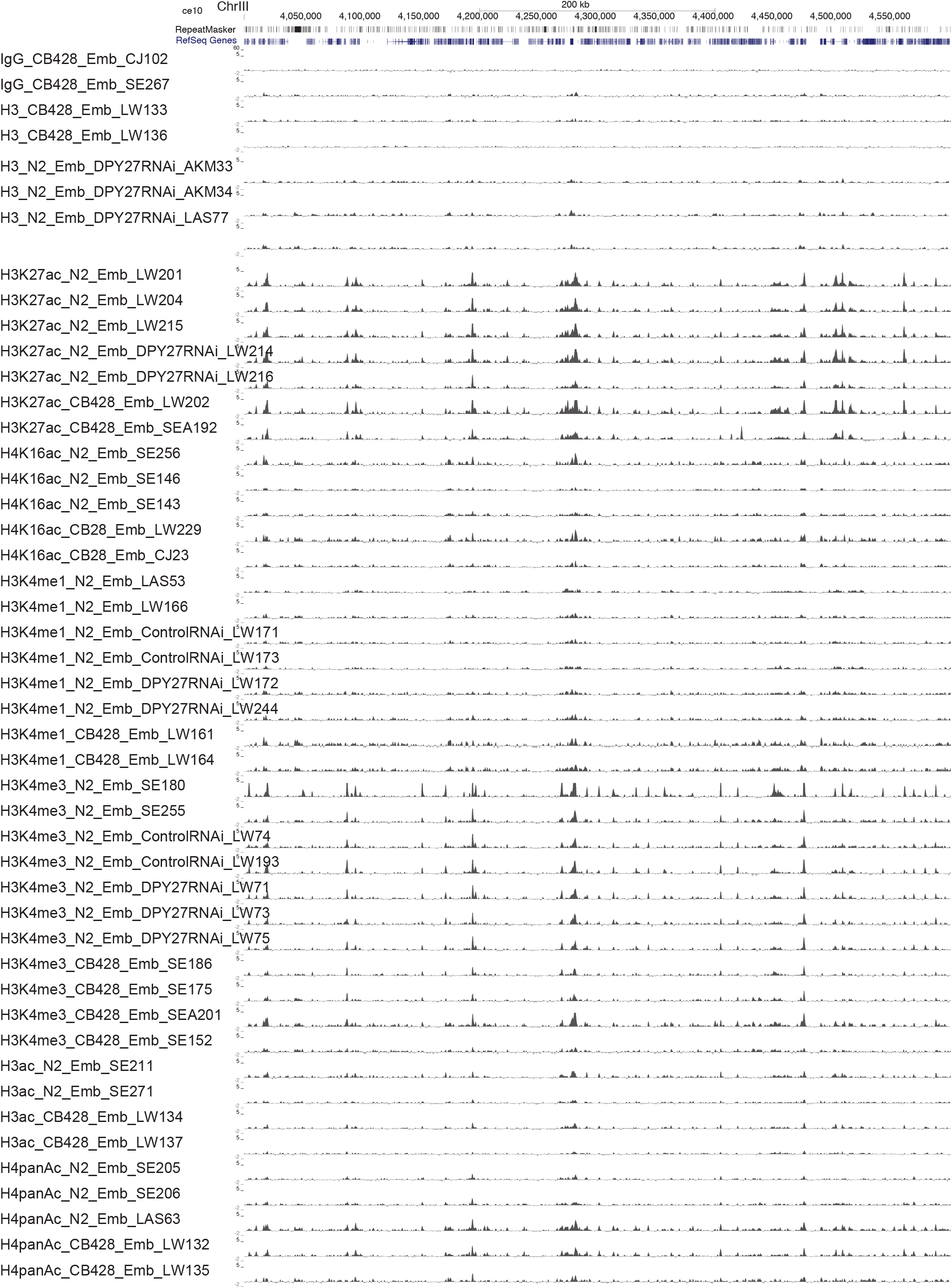
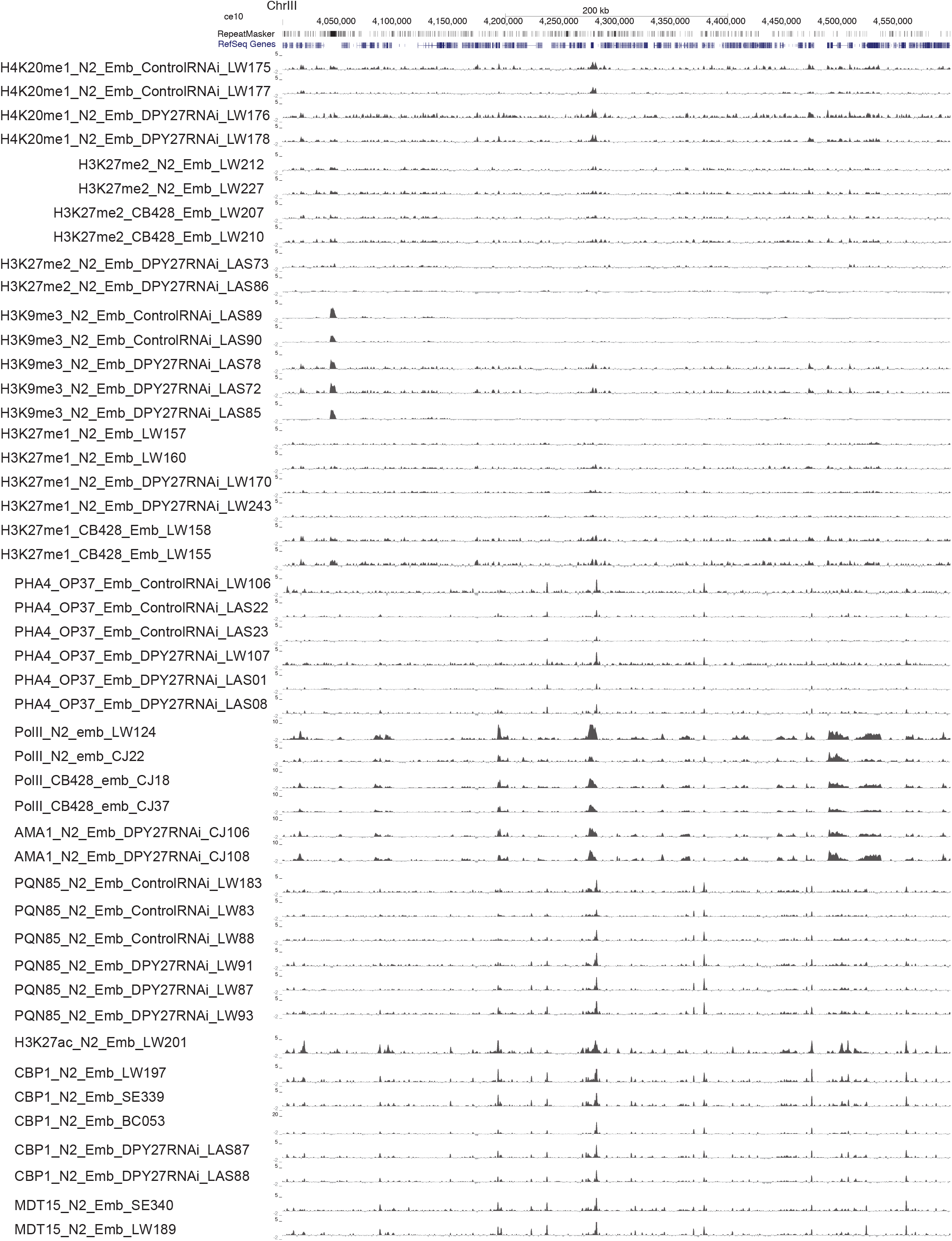
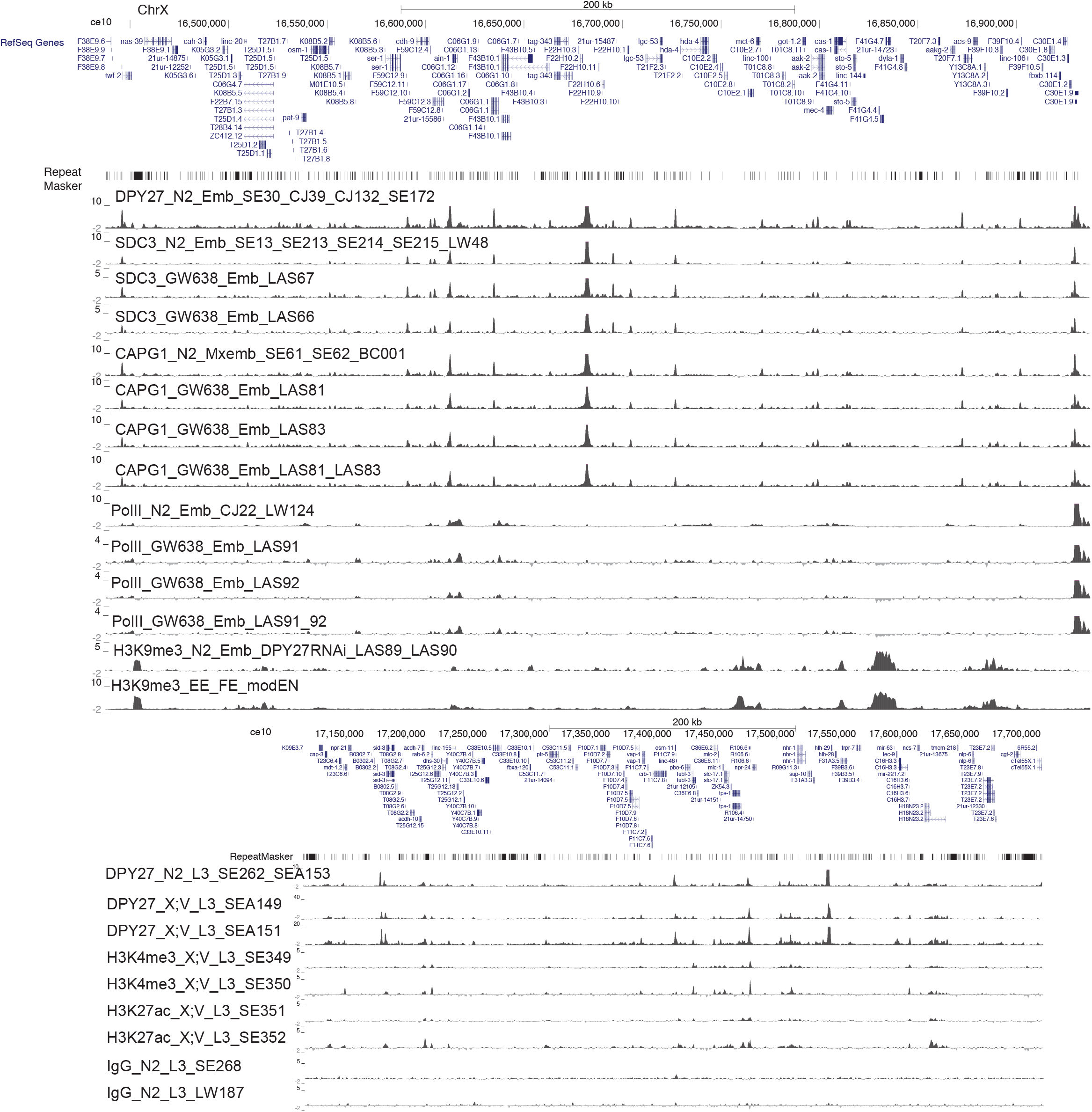
ChIP-seq binding profiles of individual biological replicates. ChIP-seq enrichment scores for each replicate are shown across a representative genomic region. The corresponding GEO entries are given in Supplemental File 1. The sample nomenclature starts with the target (e.g. H3K27ac), the strain the ChIP was performed in (e.g. N2), the developmental stage (e.g. Emb), and a unique data ID for each biological replicate (e.g. LW201)

**Supplemental Figure 2.**
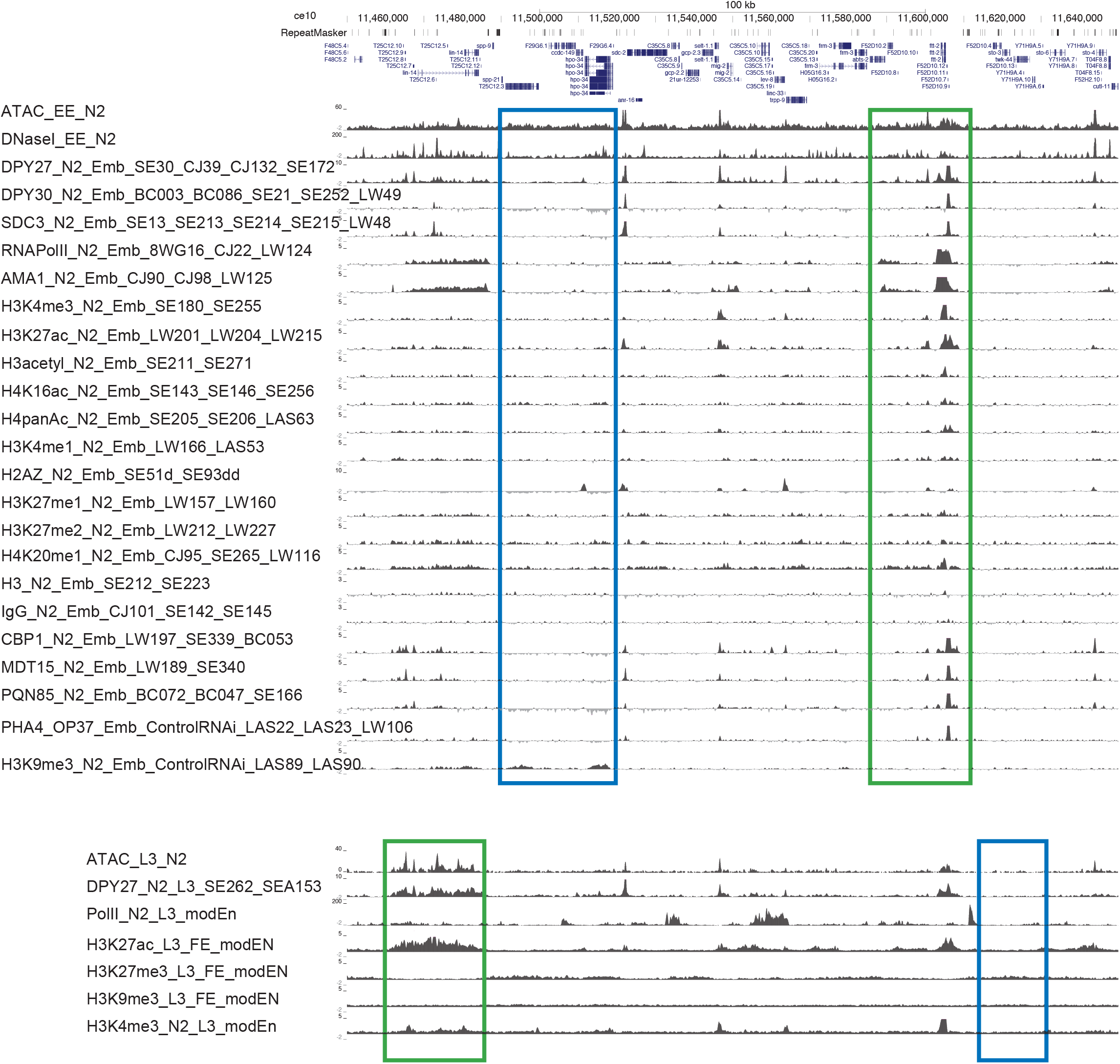
ChIP-seq ATAC-seq and DNase-seq profiles in embryo and L3 worms. ChIP-seq binding profiles for averaged data sets across a representative region of the X. Transcriptionally inactive regions are outlined in blue and active regions in green.

**Supplemental Figure 3.**
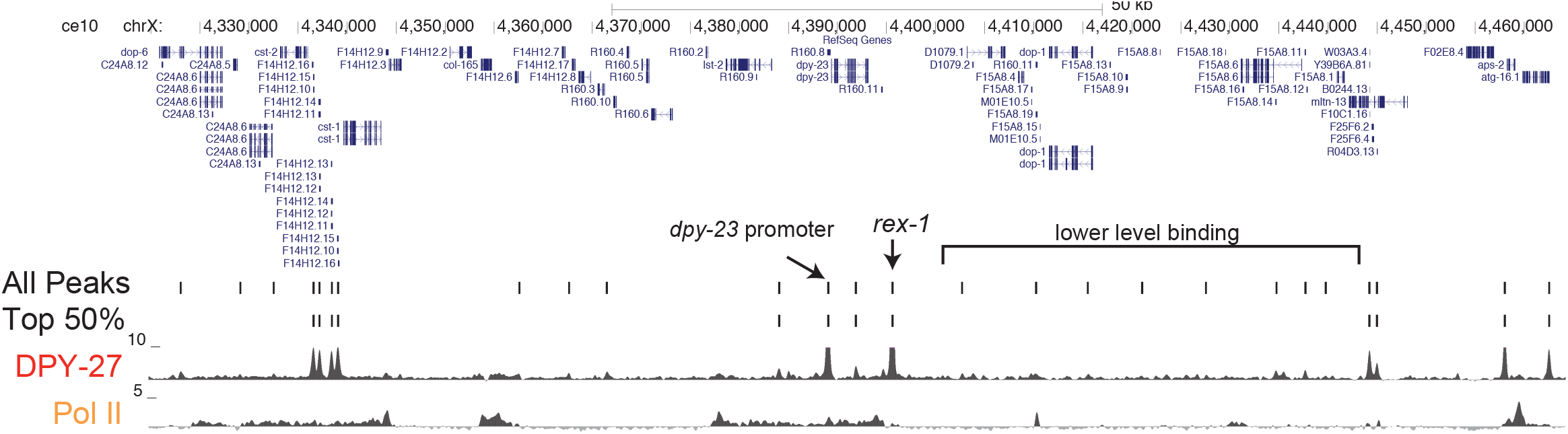
Filtering DPY-27 ChIP-seq binding peaks. DPY-27 and Pol II ChIP-seq profiles are shown along with all peaks and top 50% of peaks based on average enrichment.

**Supplemental Figure 4.**
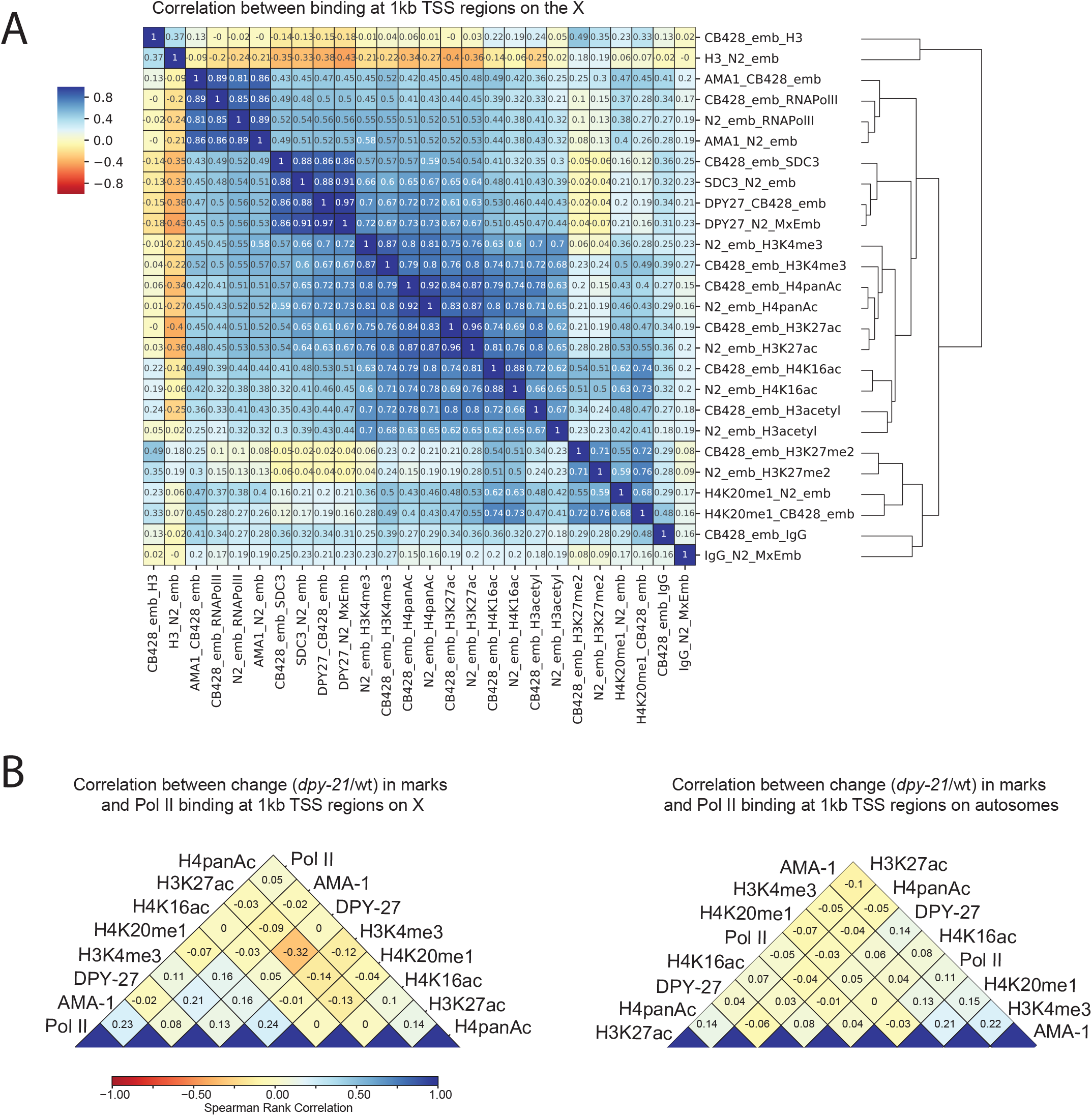
Correlation of ChIP enrichment between data sets at TSS. **(A)** Spearman rank correlations of average ChIP-seq enrichment within 1 kb regions centering at the GRO-seq defined TSS sites on the X chromosome. ChIP data from *dpy-21*, (CB428) and wild type (N2) embryos are plotted. Histone modifications associated with active transcription show positive correlation with RNA Pol II binding. Moreover, high correlation between wild type and mutant ChIP scores support the idea that the DCC does not drastically change the distribution of the marks, but rather tunes their level. **(B)** Spearman rank correlations between standardized log2 ratios of ChIP-seq enrichment (*dpy-21*,/wild type) at the 1 kb GRO-seq TSS regions across the X and autosomes. There was no strong correlation between change in RNA Pol II binding and change in histone modifications in the mutant compared to wild type. An X-specific negative correlation between DPY-27 binding and H3K27ac, supports DCC binding being linked to a reduction in H3K27ac on the X.

**Supplemental Figure 5.**
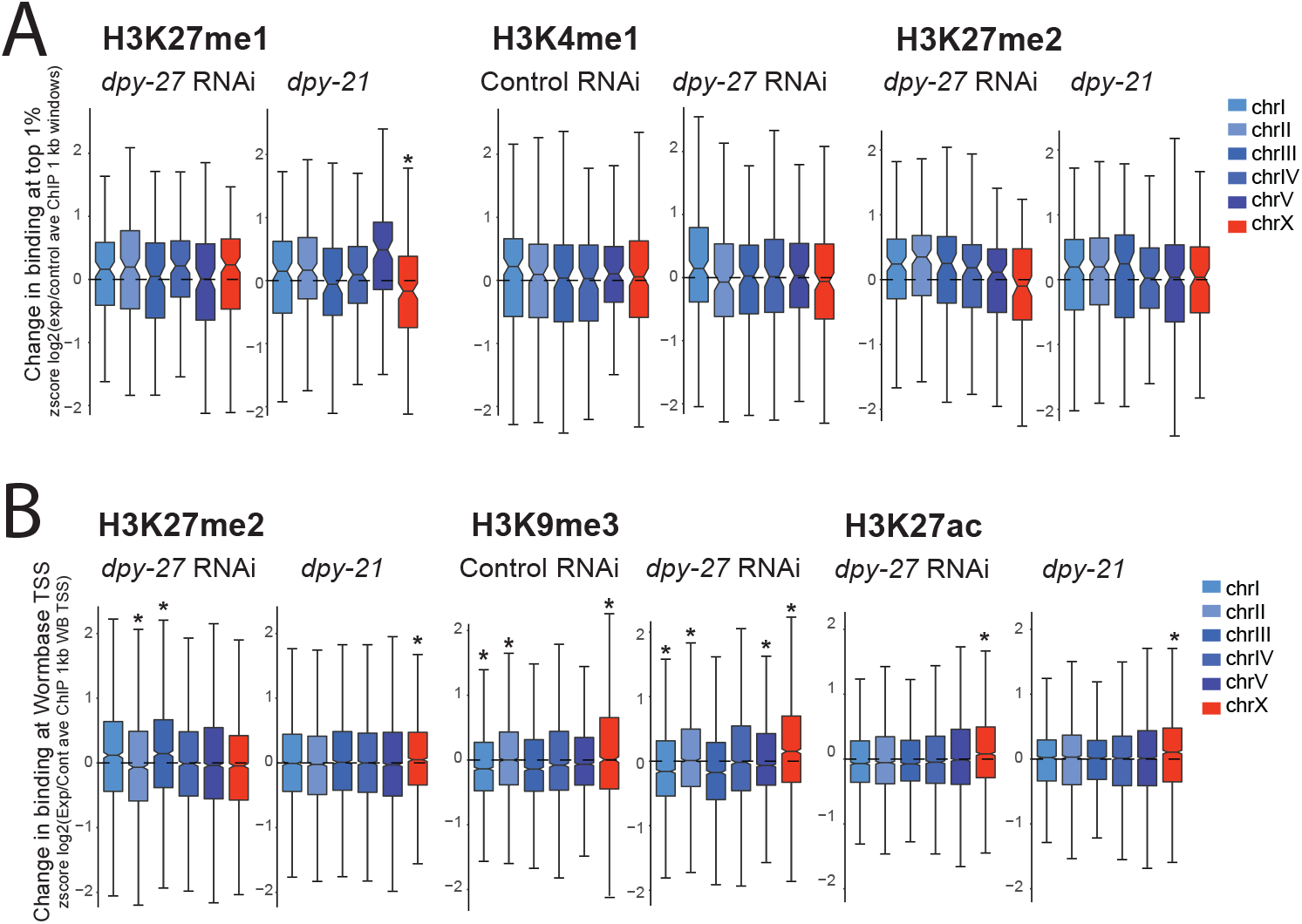
ChIP-seq enrichment changes in the DCC mutant and knockdown. **(A)** Data was analyzed as in Figure 3J and 4B, but all chromosomes are shown. DCC depletion or mutation did not significantly or specifically affect the level of H3K27me1, H3K4me1, and H3K27me2 on the X chromosomes. However, ChIP-seq data using antibodies against these modifications showed lower signal, precluding strong conclusion. **(B)** As in A, but change in histone modifications were calculated across 1 kb windows centering at Wormbase defined transcription start sites. Due to trans splicing of most genes in *C. elegans*,, these TSS coordinates are less accurate, but majority fall within 500 bp of real transcription start sites (KRUESI *et al*., 2013). DCC depletion or mutation does not specifically affect H3K27me2 and H3K9me3 but leads to increased H3K27ac across these Wormbase defined transcription start sites. Boxplots are plotted as in Figure 3. * p-value ≤ 0.001 (two-tailed Student’s t-Test).

**Supplemental Figure 6.**
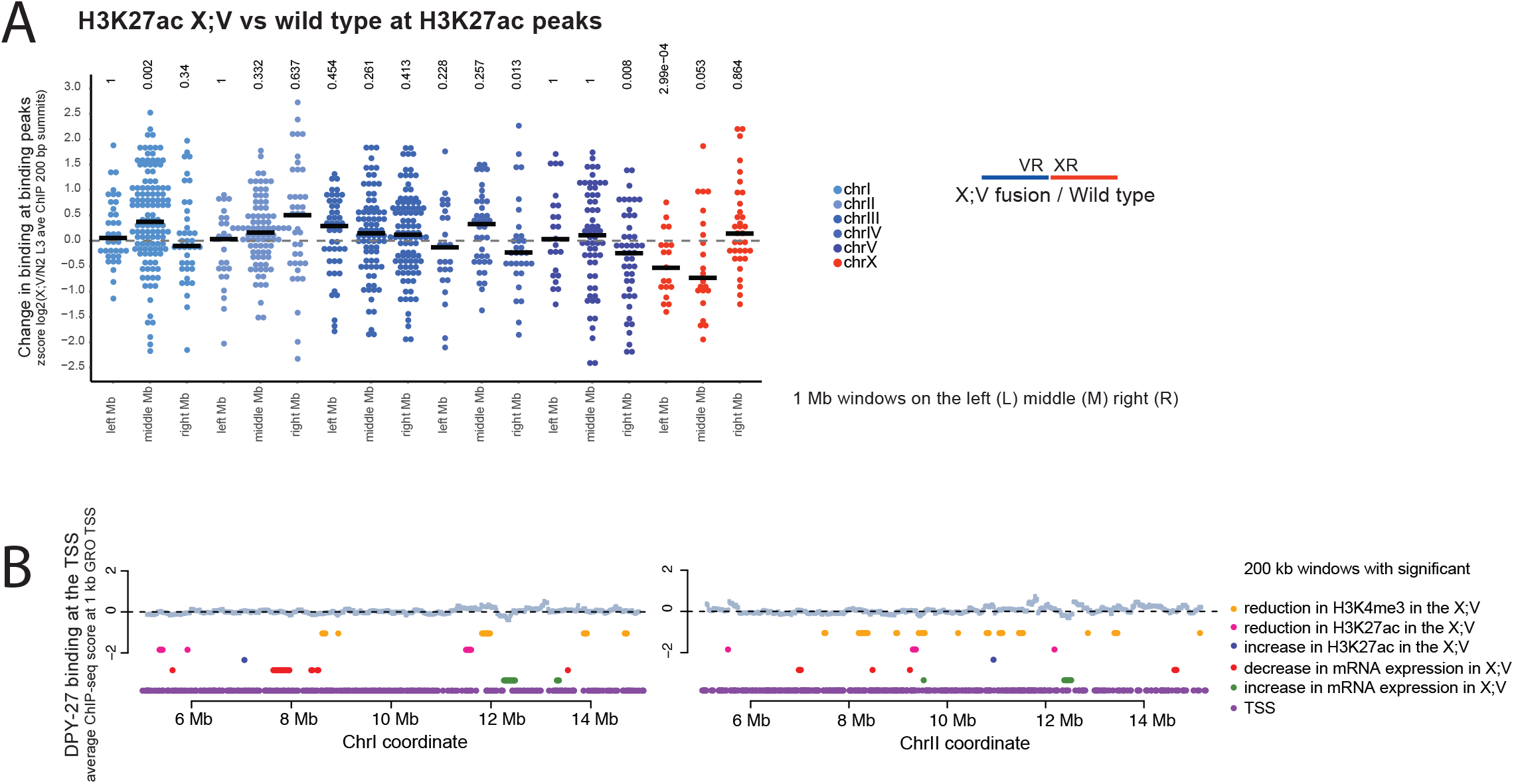
Analysis of histone modification changes in X;V fusion chromosomes. **(A)** Standardized log2 ratios of H3K27ac enrichment within 200 bp centering at the wild type H3K27ac ChIP-seq peak summits in X;V versus wild type larvae. Individual data points within the middle, left and right most 1 Mb of each chromosome are plotted. The mean ratio for each region is shown as a line. p-values were generated from a two-tailed Student’s t-Test are shown above the data. **(B)** Average DPY-27 ChIP-seq scores for 1 kb GRO-seq defined TSS regions are shown. For each 200 kb window moving along the chromosome with 20 kb steps, ChIP-seq and mRNA-seq ratios in X;V/wt were compared to the rest of the windows and a p-value statistic was generated through t-test. Windows with a = p-value ≤ 0.01 are plotted. As opposed to the chromosome V, where the windows are clustered around the region of spreading (Figure 5E), on chromosome I and II, the windows containing significant changes in gene expression and histone modifications do not show noticeable clustering.

**Supplemental Figure 7.**
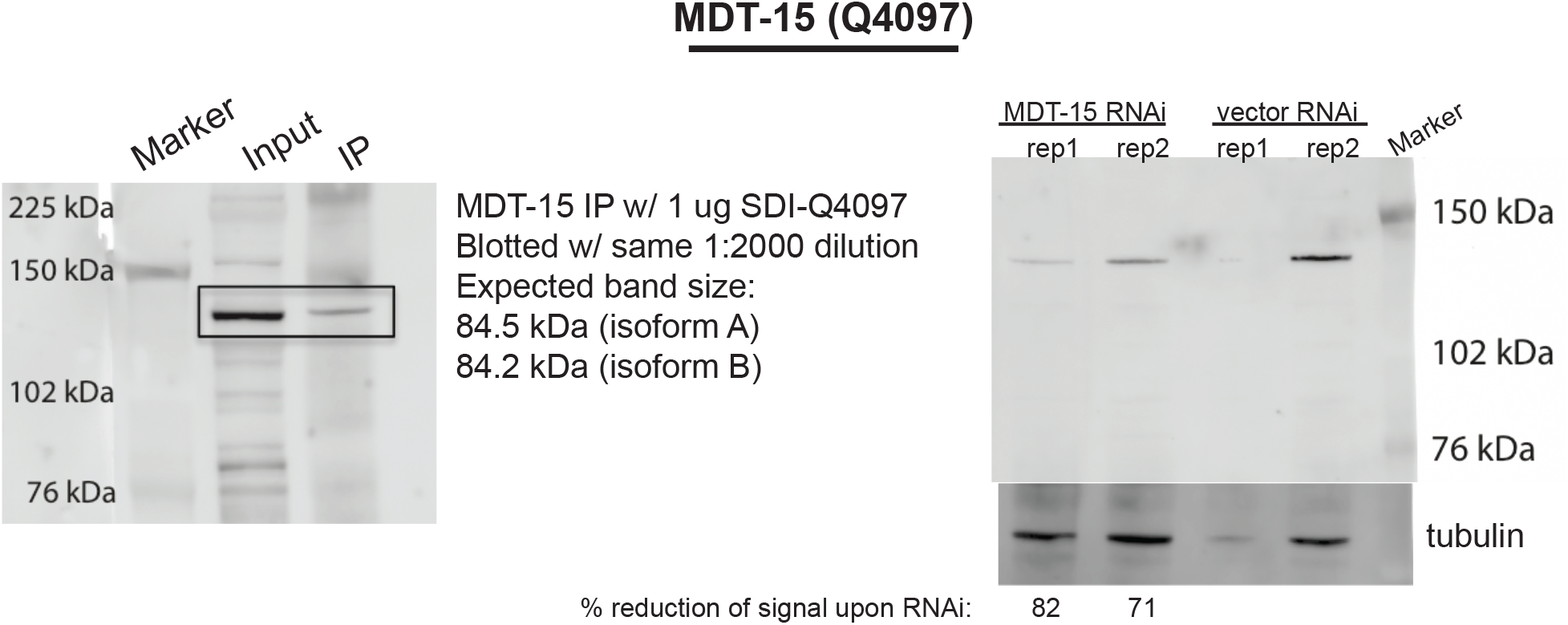
Validation of MDT-15 antibody. modENCODE generated MDT-15 Q4097 antibody was validated by western blot analysis. In N2 wild type embryos, the antibody pulled down a protein (left panel) corresponding to a band whose intensity reduced upon *mdt-15*, RNAi (compare MDT-15 signal in lane 1 to lane 4, which have similar loading based on tubulin blot). Image J was used to quantify band intensity and percentage reduction was calculated for each lane by taking a ratio of RNAi/vector MDT-15 signal versus tubulin control signal. The discrepancy between predicted MDT-15 protein size and the observed size is unclear.

## REFERENCES

Albritton, S. E., and S. Ercan, 2017 Caenorhabditis elegans Dosage Compensation: Insights into Condensin-Mediated Gene Regulation. Trends Genet.

Albritton, S. E., A. L. Kranz, P. Rao, M. Kramer, C. Dieterich et al., 2014 Sex-biased gene expression and evolution of the x chromosome in nematodes. Genetics 197: 865–883.

Albritton, S. E., A. L. Kranz, L. H. Winterkorn, L. A. Street and S. Ercan, 2017 Cooperation between a hierarchical set of recruitment sites targets the X chromosome for dosage compensation. Elife 6.

Anders, S., and W. Huber, 2010 Differential expression analysis for sequence count data. Genome Biol 11: R106.

Anders, S., P. T. Pyl and W. Huber, 2015 HTSeq--a Python framework to work with high-throughput sequencing data. Bioinformatics 31: 166–169.

Brejc, K., Q. Bian, S. Uzawa, B. S. Wheeler, E. C. Anderson et al., 2017 Dynamic Control of X Chromosome Conformation and Repression by a Histone H4K20 Demethylase. Cell.

Cano-Rodriguez, D., R. A. Gjaltema, L. J. Jilderda, P. Jellema, J. Dokter-Fokkens et al., 2016 Writing of H3K4Me3 overcomes epigenetic silencing in a sustained but context-dependent manner. Nat Commun 7: 12284.

Crane, E., Q. Bian, R. P. McCord, B. R. Lajoie, B. S. Wheeler et al., 2015 Condensin-driven remodelling of X chromosome topology during dosage compensation. Nature 523: 240–244.

Csankovszki, G., K. Collette, K. Spahl, J. Carey, M. Snyder et al., 2009 Three distinct condensin complexes control C. elegans chromosome dynamics. Curr Biol 19: 9–19.

Csankovszki, G., P. McDonel and B. J. Meyer, 2004 Recruitment and spreading of the C. elegans dosage compensation complex along X chromosomes. Science 303: 1182–1185.

Cuylen, S., J. Metz, A. Hruby and C. H. Haering, 2013 Entrapment of chromosomes by condensin rings prevents their breakage during cytokinesis. Dev Cell 27: 469–478.

D’Ambrosio, C., C. K. Schmidt, Y. Katou, G. Kelly, T. Itoh et al., 2008 Identification of cis-acting sites for condensin loading onto budding yeast chromosomes. Genes Dev 22: 2215–2227.

Daugherty, A. C., R. W. Yeo, J. D. Buenrostro, W. J. Greenleaf, A. Kundaje et al., 2017 Chromatin accessibility dynamics reveal novel functional enhancers in C. elegans. Genome Res 27: 2096–2107.

Dowen, J. M., and R. A. Young, 2014 SMC complexes link gene expression and genome architecture. Curr Opin Genet Dev 25: 131–137.

Ercan, S., L. L. Dick and J. D. Lieb, 2009 The C. elegans dosage compensation complex propagates dynamically and independently of X chromosome sequence. Curr Biol 19: 1777–1787.

Ercan, S., P. G. Giresi, C. M. Whittle, X. Zhang, R. D. Green et al., 2007 X chromosome repression by localization of the C. elegans dosage compensation machinery to sites of transcription initiation. Nat Genet 39: 403–408.

Gerstein, M. B., Z. J. Lu, E. L. Van Nostrand, C. Cheng, B. I. Arshinoff et al., 2010 Integrative analysis of the Caenorhabditis elegans genome by the modENCODE project. Science 330: 1775–1787.

Ginno, P. A., L. Burger, J. Seebacher, V. Iesmantavicius and D. Schubeler, 2018 Cell cycle-resolved chromatin proteomics reveals the extent of mitotic preservation of the genomic regulatory landscape. Nat Commun 9: 4048.

Haeusler, R. A., M. Pratt-Hyatt, P. D. Good, T. A. Gipson and D. R. Engelke, 2008 Clustering of yeast tRNA genes is mediated by specific association of condensin with tRNA gene transcription complexes. Genes Dev 22: 2204–2214.

Hirano, T., 2006 At the heart of the chromosome: SMC proteins in action. Nat Rev Mol Cell Biol 7: 311–322.

Hirano, T., 2016 Condensin-Based Chromosome Organization from Bacteria to Vertebrates. Cell 164: 847–857.

Ho, M. C. W., P. Quintero-Cadena and P. W. Sternberg, 2017 Genome-wide discovery of active regulatory elements and transcription factor footprints in Caenorhabditis elegans using DNase-seq. Genome Res 27: 2108–2119.

Howe, F. S., H. Fischl, S. C. Murray and J. Mellor, 2017 Is H3K4me3 instructive for transcription activation? Bioessays 39: 1–12.

Iwasaki, O., A. Tanaka, H. Tanizawa, S. I. Grewal and K. Noma, 2010 Centromeric localization of dispersed Pol III genes in fission yeast. Mol Biol Cell 21: 254–265.

Iwasaki, O., H. Tanizawa, K. D. Kim, Y. Yokoyama, C. J. Corcoran et al., 2015 Interaction between TBP and Condensin Drives the Organization and Faithful Segregation of Mitotic Chromosomes. Mol Cell 59: 755–767.

Jans, J., J. M. Gladden, E. J. Ralston, C. S. Pickle, A. H. Michel et al., 2009 A condensin-like dosage compensation complex acts at a distance to control expression throughout the genome. Genes Dev 23: 602–618.

Jeppsson, K., T. Kanno, K. Shirahige and C. Sjogren, 2014 The maintenance of chromosome structure: positioning and functioning of SMC complexes. Nat Rev Mol Cell Biol 15: 601–614.

Karlic, R., H. R. Chung, J. Lasserre, K. Vlahovicek and M. Vingron, 2010 Histone modification levels are predictive for gene expression. Proc Natl Acad Sci U S A 107: 2926–2931.

Kim, K. D., H. Tanizawa, O. Iwasaki and K. I. Noma, 2016 Transcription factors mediate condensin recruitment and global chromosomal organization in fission yeast. Nat Genet.

Kramer, M., A. L. Kranz, A. Su, L. H. Winterkorn, S. E. Albritton et al., 2015 Developmental Dynamics of X-Chromosome Dosage Compensation by the DCC and H4K20me1 in C. elegans. PLoS Genet 11: e1005698.

Kramer, M., P. Rao and S. Ercan, 2016 Untangling the Contributions of Sex-Specific Gene Regulation and X-Chromosome Dosage to Sex-Biased Gene Expression in Caenorhabditis elegans. Genetics 204: 355–369.

Kranz, A. L., C. Y. Jiao, L. H. Winterkorn, S. E. Albritton, M. Kramer et al., 2013 Genome-wide analysis of condensin binding in Caenorhabditis elegans. Genome Biol 14: R112.

Kruesi, W. S., L. J. Core, C. T. Waters, J. T. Lis and B. J. Meyer, 2013 Condensin controls recruitment of RNA polymerase II to achieve nematode X-chromosome dosage compensation. Elife 2: e00808.

Kschonsak, M., F. Merkel, S. Bisht, J. Metz, V. Rybin et al., 2017 Structural Basis for a Safety-Belt Mechanism That Anchors Condensin to Chromosomes. Cell 171: 588–600 e524.

Langmead, B., C. Trapnell, M. Pop and S. L. Salzberg, 2009 Ultrafast and memory-efficient alignment of short DNA sequences to the human genome. Genome Biol 10: R25.

Lau, A. C., K. Nabeshima and G. Csankovszki, 2014 The C. elegans dosage compensation complex mediates interphase X chromosome compaction. Epigenetics Chromatin 7: 31.

Leatham-Jensen, M., C. M. Uyehara, B. D. Strahl, A. G. Matera, R. J. Duronio et al., 2019 Lysine 27 of replication-independent histone H3.3 is required for Polycomb target gene silencing but not for gene activation. PLoS Genet 15: e1007932.

Lowden, M. R., B. Meier, T. W. Lee, J. Hall and S. Ahmed, 2008 End joining at Caenorhabditis elegans telomeres. Genetics 180: 741–754.

Maxwell, C. S., W. S. Kruesi, L. J. Core, N. Kurhanewicz, C. T. Waters et al., 2014 Pol II docking and pausing at growth and stress genes in C. elegans. Cell Rep 6: 455–466.

McDonel, P., J. Jans, B. K. Peterson and B. J. Meyer, 2006 Clustered DNA motifs mark X chromosomes for repression by a dosage compensation complex. Nature 444: 614–618.

McKay, D. J., S. Klusza, T. J. Penke, M. P. Meers, K. P. Curry et al., 2015 Interrogating the function of metazoan histones using engineered gene clusters. Dev Cell 32: 373–386.

Paul, M. R., A. Hochwagen and S. Ercan, 2018 Condensin action and compaction. Curr Genet.

Perez-Lluch, S., E. Blanco, H. Tilgner, J. Curado, M. Ruiz-Romero et al., 2015 Absence of canonical marks of active chromatin in developmentally regulated genes. Nat Genet 47: 1158–1167.

Pferdehirt, R. R., W. S. Kruesi and B. J. Meyer, 2011 An MLL/COMPASS subunit functions in the C. elegans dosage compensation complex to target X chromosomes for transcriptional regulation of gene expression. Genes Dev 25: 499–515.

Ramirez, F., F. Dundar, S. Diehl, B. A. Gruning and T. Manke, 2014 deepTools: a flexible platform for exploring deep-sequencing data. Nucleic Acids Res 42: W187–191.

Robellet, X., V. Vanoosthuyse and P. Bernard, 2017 The loading of condensin in the context of chromatin. Curr Genet 63: 577–589.

Rowley, M. J., and V. G. Corces, 2018 Organizational principles of 3D genome architecture. Nat Rev Genet.

Salmon-Divon, M., H. Dvinge, K. Tammoja and P. Bertone, 2010 PeakAnalyzer: genome-wide annotation of chromatin binding and modification loci. BMC Bioinformatics 11: 415.

Sharma, R., D. Jost, J. Kind, G. Gomez-Saldivar, B. van Steensel et al., 2014 Differential spatial and structural organization of the X chromosome underlies dosage compensation in C. elegans. Genes Dev 28: 2591–2596.

Snyder, M. J., A. C. Lau, E. A. Brouhard, M. B. Davis, J. Jiang et al., 2016 Anchoring of Heterochromatin to the Nuclear Lamina Reinforces Dosage Compensation-Mediated Gene Repression. PLoS Genet 12: e1006341.

Stasevich, T. J., Y. Hayashi-Takanaka, Y. Sato, K. Maehara, Y. Ohkawa et al., 2014 Regulation of RNA polymerase II activation by histone acetylation in single living cells. Nature 516: 272–275.

Sutani, T., T. Sakata, R. Nakato, K. Masuda, M. Ishibashi et al., 2015 Condensin targets and reduces unwound DNA structures associated with transcription in mitotic chromosome condensation. Nat Commun 6: 7815.

Towbin, B. D., C. Gonzalez-Aguilera, R. Sack, D. Gaidatzis, V. Kalck et al., 2012 Step-wise methylation of histone H3K9 positions heterochromatin at the nuclear periphery. Cell 150: 934–947.

Trapnell, C., A. Roberts, L. Goff, G. Pertea, D. Kim et al., 2012 Differential gene and transcript expression analysis of RNA-seq experiments with TopHat and Cufflinks. Nature Protocols 7: 562–578.

Van Bortle, K., M. H. Nichols, L. Li, C. T. Ong, N. Takenaka et al., 2014 Insulator function and topological domain border strength scale with architectural protein occupancy. Genome Biol 15: R82.

van Ruiten, M. S., and B. D. Rowland, 2018 SMC Complexes: Universal DNA Looping Machines with Distinct Regulators. Trends Genet.

Vielle, A., J. Lang, Y. Dong, S. Ercan, C. Kotwaliwale et al., 2012 H4K20me1 contributes to downregulation of X-linked genes for C. elegans dosage compensation. PLoS Genet 8: e1002933.

Wells, M. B., M. J. Snyder, L. M. Custer and G. Csankovszki, 2012 Caenorhabditis elegans dosage compensation regulates histone H4 chromatin state on X chromosomes. Mol Cell Biol 32: 1710–1719.

Yuen, K. C., B. D. Slaughter and J. L. Gerton, 2017 Condensin II is anchored by TFIIIC and H3K4me3 in the mammalian genome and supports the expression of active dense gene clusters. Sci Adv 3: e1700191.

Zhang, Y., T. Liu, C. A. Meyer, J. Eeckhoute, D. S. Johnson et al., 2008 Model-based analysis of ChIP-Seq (MACS). Genome Biol 9: R137.

Zhang, Z., and M. Q. Zhang, 2011 Histone modification profiles are predictive for tissue/cell-type specific expression of both protein-coding and microRNA genes. BMC Bioinformatics 12: 155.

